# Denoising task-correlated head motion from motor-task fMRI data with multi-echo ICA

**DOI:** 10.1101/2023.07.19.549746

**Authors:** Neha A. Reddy, Kristina M. Zvolanek, Stefano Moia, César Caballero-Gaudes, Molly G. Bright

**Affiliations:** Department of Physical Therapy and Human Movement Sciences, Feinberg School of Medicine, Northwestern University, Chicago, IL, United States; Department of Biomedical Engineering, McCormick School of Engineering and Applied Sciences, Northwestern University, Evanston, IL, United States; Basque Center on Cognition, Brain and Language, Donostia, Gipuzkoa, Spain; Neuro-X Institute, École polytechnique fédérale de Lausanne, Geneva, Switzerland; Department of Radiology and Medical Informatics (DRIM), Faculty of Medicine, University of Geneva, Geneva, Switzerland

**Keywords:** BOLD fMRI, motor task, task-correlated head motion, multi-echo, independent component analysis

## Abstract

Motor-task functional magnetic resonance imaging (fMRI) is crucial in the study of several clinical conditions, including stroke and Parkinson’s disease. However, motor-task fMRI is complicated by task-correlated head motion, which can be magnified in clinical populations and confounds motor activation results. One method that may mitigate this issue is multi-echo independent component analysis (ME-ICA), which has been shown to separate the effects of head motion from the desired BOLD signal but has not been tested in motor-task datasets with high amounts of motion. In this study, we collected an fMRI dataset from a healthy population who performed a hand grasp task with and without task-correlated amplified head motion to simulate a motor-impaired population. We analyzed these data using three models: single-echo (SE), multi-echo optimally combined (ME-OC), and ME-ICA. We compared the models’ performance in mitigating the effects of head motion on the subject level and group level. On the subject level, ME-ICA better dissociated the effects of head motion from the BOLD signal and reduced noise. Both ME models led to increased t-statistics in brain motor regions. In scans with high levels of motion, ME-ICA additionally mitigated artifacts and increased stability of beta coefficient estimates, compared to SE. On the group level, all three models produced activation clusters in expected motor areas in scans with both low and high motion, indicating that group-level averaging may also sufficiently resolve motion artifacts that vary by subject. These findings demonstrate that ME-ICA is a useful tool for subject-level analysis of motor-task data with high levels of task-correlated head motion. The improvements afforded by ME-ICA are critical to improve reliability of subject-level activation maps for clinical populations in which group-level analysis may not be feasible or appropriate, for example in a chronic stroke cohort with varying stroke location and degree of tissue damage.

## 1. Introduction

Assessing motor function with functional magnetic resonance imaging (fMRI) is critical in several clinical and research applications, including surgical planning (Ciavarro et al., 2021; Heilbrun et al., 2001; Vysotski et al., 2018; Wilkinson et al., 2003), tracking clinical outcomes after interventions (Péran et al., 2020; Stephan et al., 2001; Ward et al., 2003a, 2003b), the study of motor neurophysiology and rehabilitation in stroke and cerebral palsy (Araneda et al., 2021; Cramer, Mark, et al., 2002; Hannanu et al., 2020; Hermsdörfer et al., 2003; J.P. et al., 2007), and the study of Parkinson’s disease (Martin et al., 2019; Sabatini et al., 1998; Wu & Hallett, 2005). However, head motion poses a serious problem during motor-task fMRI, since motion confounds interpretation of fMRI results by causing false positive and negative activation in task-fMRI (Friston et al., 1996) and spurious correlations in functional connectivity data (Power et al., 2012; Satterthwaite et al., 2012; van Dijk et al., 2012). In particular, certain clinical populations may have amplified head motion during motor actions; for example, stroke participants were observed to have twice the amount of head motion during hand grasp and ankle flexion tasks compared to healthy individuals (Seto et al., 2001). Head motion is also increased in participants who may be uncomfortable in the MRI environment, such as children (Byars et al., 2002; Kotsoni et al., 2006).

Several strategies have been employed to address the issue of motion correction for fMRI data (Caballero-Gaudes & Reynolds, 2017). Most commonly, volume registration is performed to align images across the scan time points; however, it has been shown that this technique is not sufficient to eliminate motion effects (Freire & Mangin, 2001; Grootoonk et al., 2000). Many studies take a further step of adding the motion parameters and their derivatives calculated during volume registration as regressors during modeling (Friston et al., 1996), though motion-related confounds may still be present in the data and result in spurious findings (Power et al., 2012), particularly when the motion is task-correlated (Johnstone et al., 2006). Censoring and interpolation are approaches that remove large spikes in motion above a set threshold, but drawbacks are disruption of a signal’s temporal correlation and challenges in deciding the correct threshold (Caballero-Gaudes & Reynolds, 2017). Additionally, censoring and interpolation do not correct for smaller motions throughout the scan, which may be more prevalent in clinical populations and can also lead to false results (Power et al., 2012). Other commonly used tools to separate effects of noise from the desired Blood Oxygenation Level Dependent (BOLD) signal include approaches based on Principal Component Analysis (PCA), such as CompCor (Behzadi et al., 2007; Muschelli et al., 2014) and GLMdenoise (Kay et al., 2013) and approaches based on Independent Component Analysis (ICA), such as FIX-ICA (Griffanti et al., 2014; Salimi-Khorshidi et al., 2014) and ICA-AROMA (Pruim et al., 2015), that identify noise components of the signal and regress out their corresponding timecourses.

Additional challenges arise when head motion is correlated with the task stimulus (Field et al., 2000; Hajnal et al., 1994; Johnstone et al., 2006; Moia et al., 2021; Soltysik & Hyde, 2006; Xu et al., 2014) and the impact of task-correlated motion on fMRI activation results is not easily corrected for (Mumford et al., 2015). For example, some of the motion correction strategies described above have been shown to lose efficacy when motion is task-correlated; such strategies include volume realignment (Field et al., 2000; Morgan et al., 2007), adding motion regressors to the model (Johnstone et al., 2006), and PCA (Patriat et al., 2017). Of note, motor tasks have been associated with increased task-correlated head motion in a motor-impaired population (Seto et al., 2001), suggesting that this issue may be amplified in certain clinical cohorts.

An efficient method to identify task-correlated confounds in fMRI data involves separating the effects of changes in the transverse relaxation parameter T_2_*, which are related to the BOLD response, from the effects of changes in the net magnetization S_0_, which include motion, pulsation, and inflow effects (Menon et al., 1993; Ogawa et al., 1993). This approach builds on the physical principles of the T_2_*-weighted fMRI signal and the BOLD response. In initial studies, a short echo time (TE) closely following the radiofrequency (RF) excitation was used to approximate S_0_ effects and remove them from the T_2_*-related BOLD fMRI signal. Several studies used dual-echo (Glover et al. 1996; Ing and Schwarzbauer 2012; Bright and Murphy 2013; Buur, Poser, and Norris 2009) and multi-echo (Barth et al. 1999) acquisitions to perform this noise separation technique in fMRI data. Kundu et al. (2012, 2013) proposed a related method to separate BOLD and non-BOLD components using multi-echo information and ICA (ME-ICA) to calculate two parameters, kappa and rho, that quantify the T_2_*- and S_0_-weighting of the components, respectively. Along with other features, these kappa and rho parameters can be used to understand the likeliness that each component is attributed to the true BOLD signal, and subsequently better classify the component (Dupre et al., 2021). ME-ICA has been used with success in various resting-state (Cohen, Chang, et al., 2021; Cohen, Yang, et al., 2021; Dipasquale et al., 2017) and task-based MRI (Cohen et al., 2018; Cohen, Jagra, Visser, et al., 2021; Cohen, Jagra, Yang, et al., 2021; Cohen & Wang, 2019; Evans et al., 2015; Gonzalez-Castillo et al., 2016; Lombardo et al., 2016; Moia et al., 2021) applications.

However, studies have not yet evaluated the use of ME-ICA in denoising task-correlated head motion in motor-task fMRI data, where it could be critically useful for the clinical applications described previously, and other correction methods may fail. Motor areas in healthy individuals have been identified in cortical (Gordon et al., 2023) and cerebellar (Ashida et al., 2019; Spencer et al., 2007; Stoodley, 2012) regions. Multi-echo information might be additionally useful for improving our detection in non-cortical regions implicated in human movement, such as the cerebellum: several studies have observed that multi-echo and ME-ICA methods lead to signal improvements in subcortical regions (Gonzalez-Castillo et al., 2016; Guediche et al., 2021; Lombardo et al., 2016; Lynch et al., 2020).

In this study, we apply ME-ICA to a motor-task dataset including a wide range of task-correlated head motion amplitudes, to evaluate its utility in improving the interpretability of motor-task fMRI in clinical populations. To focus on the effects of task-correlated head motion, distinct from other neural or vascular confounds present in clinical populations, we simulate amplified task-correlated head motion in healthy participants as done in previous studies (Bright & Murphy, 2013; Buur et al., 2009; Ing & Schwarzbauer, 2012; Kochiyama et al., 2005). We evaluate the motion characteristics of this dataset, including average head motion and task-correlation of motion, and compare these characteristics to those of two representative stroke participants. Then, we probe the effects of ME and ME-ICA analysis on subject- and group-level activation results, including parameter estimates and t-statistics in relevant brain motor areas.

## 2. Methods

### 2.1. Data Collection

This study was approved by the Northwestern University Institutional Review Board, and all participants provided written, informed consent. Eight right-handed, healthy participants with no history of neurological or vascular disorders (4M, 26 ± 2 years) were scanned on a Siemens 3T Prisma MRI system with a 64-channel head coil. A structural T1-weighted multi-echo MPRAGE image was collected using parameters adapted from Tisdall and colleagues (2016): TR=2.17 s, TEs=1.69/3.55/5.41 ms, TI=1.16 s, FA=7°, FOV=256×256 mm^2^, and voxel size=1×1×1 mm^3^. The three echo MPRAGE images were combined using root-mean-square. Two functional scans were collected using a multi-band multi-echo gradient-echo echo planar imaging sequence provided by the Center for Magnetic Resonance Research (CMRR, Minnesota): TR=2 s, TEs=10.8/28.03/45.26/62.49/79.72 ms, FA=70°, MB factor=4, GRAPPA=2, voxel size=2.5×2.5×2 mm^3^, 210 volumes (Moeller et al., 2010; Setsompop et al., 2012). Tape placed across the forehead provided tactile feedback of head motion, as found to reduce translational and rotational motion (Krause et al., 2019). During the scans, CO_2_ was measured via a nasal cannula and gas analyzer at a sampling rate of 20 Hz (PowerLab, ADInstruments).

#### 2.1.1. Motor task

Hand grasp tasks were performed with a device that interfaced with the scanner bed and contained two load cells (Interface Inc., Scottsdale, AZ, USA). This device provided support to both arms, positioning them at an elbow flexion of approximately 30 degrees, and provided a comfortable grasping interface with the load cells for each hand. Prior to the scan, grasp forces at maximum voluntary contraction (MVC) were calculated for the right and left hand as the average of three unilateral maximum grasp force trials. During the task trials, participants performed an isometric unimanual grasping task at 40% MVC. Each task trial was a 10-s “squeeze” and 15-s “relax”, alternating 4 trials per hand, for a total of 16 trials. Four participants began with the right-hand grasp, and four participants began with the left hand. Task instructions and real-time force feedback from the load cells were viewed on a screen to facilitate force targeting. Text on the screen instructed the start and end of the hand grasp periods. During the hand grasp periods, a target box of 35-45% MVC indicated the target force level. A moving bar indicated the participants’ real-time force level and turned green when the force was within the target range.

During the first functional scan *(Limited)*, participants were instructed to keep their head as still as possible. During the second functional scan *(Amplified)*, participants were instructed to add a small downwards nod at the beginning of each hand grasp and a small upwards nod to return to the starting position upon release of each hand grasp; this instruction added self-directed task-correlated motion to the Amplified motion scan. Out of the six possible directions of head rotation and translation, nodding was chosen as it was a primary direction of motion observed in preliminary data in healthy and stroke participants, as well as feasible to be performed within the limitations of the MRI environment. Right- and left-hand grasp force data from the load cells were measured continuously during the scans at a sampling rate of 20 Hz and synchronized with CO_2_ recordings and the scanner trigger (PowerLab, ADInstruments).

### 2.2. Data Analysis

Data are publicly available on OpenNeuro at doi:10.18112/openneuro.ds004662.v1.1.0 (Reddy et al., 2023). Code is publicly available at https://github.com/BrightLab-ANVIL/PreProc_BRAIN and https://github.com/BrightLab-ANVIL/Reddy_MotorMEICA.

#### 2.2.1. Creation of motor task and end-tidal CO_2_ regressors

Using MATLAB (version 2020a), the force traces from the right and left hands during the scans were normalized to the participant’s maximum grip force, then convolved with a canonical hemodynamic response function. The resulting traces were rescaled to the range of the normalized force traces, then demeaned. The demeaned force trace was then downsampled to the resolution of the functional MRI images (TR=2s) in order to generate corresponding RGrip and LGrip regressors.

End-tidal peaks were detected in the CO_2_ data using an automatic peak-finder in MATLAB, then manually inspected. The end-tidal peaks were interpolated to form an end-tidal CO_2_ trace that was convolved with a canonical hemodynamic response function, rescaled to the range of the unconvolved timeseries, then demeaned. The demeaned trace was then downsampled to the resolution of the functional MRI images (TR=2s).

#### 2.2.2. Structural MRI pre-processing

T1-weighted images for each subject were processed with FSL’s (Jenkinson et al., 2012) fsl_anat, which performs bias field correction and brain extraction.

#### 2.2.3. Functional MRI pre-processing

FSL (version 6.0.7.2) (Jenkinson et al., 2012) and AFNI (version 23.2.12) (Cox J.S., 1996) tools were used for fMRI preprocessing of single-echo and multi-echo datasets as follows.

##### Single-echo (SE)

The second echo data (TE=28.03 ms) were used as a proxy for conventional single-echo (SE) fMRI analysis. The first 10 volumes of each scan were removed to allow the signal to achieve steady-state magnetization. Head-motion realignment was computed with reference to the Single Band reference image taken at the start of the scan (3dvolreg, AFNI). After brain extraction (bet, FSL), the SE timeseries were converted to signal percentage change for further analysis. No spatial smoothing was performed.

##### Multi-echo (ME)

The first 10 volumes of each echo were removed to allow the signal to achieve steady-state magnetization. Head-motion realignment was estimated for the first echo with reference to the Single Band reference image taken at the start of the scan (3dvolreg, AFNI) and then applied to all echoes (3dAllineate, AFNI). All images were brain extracted (bet, FSL). Tedana (version 0.0.11) (Ahmed et al., 2021; Dupre et al., 2021) was used to calculate a T_2_*-weighted combination of the five echo datasets, producing the optimally combined (ME-OC) fMRI dataset. The voxel-wise T_2_* estimates calculated by tedana for use in optimal combination of echoes are shown in **Supplemental Figure 1**. The ME-OC timeseries was converted to signal percentage change for further analysis. No spatial smoothing was performed.

As tedana leads to a tighter brain mask through removal of additional voxels with insufficient signal, for the subject-level analyses described subsequently, this tighter brain mask is applied to both SE and ME analysis for more appropriate comparison between models.

##### Multi-echo Independent Component Analysis (ME-ICA)

Multi-echo independent component analysis (ME-ICA) was performed on the ME-OC fMRI data (prior to conversion to signal percentage change) using tedana (version 0.0.11). The resulting components were manually classified (accepted as signal of interest or rejected as noise), using criteria adapted from (Griffanti et al., 2017), and aided by Rica (Uruñuela, 2021). Components were *rejected* if any of the following applied:

1. Spatial maps showing

a. Larger amplitudes mostly in white matter or CSF voxels;
b. Small and scattered clusters
c. Striped alternating positive and negative regions (indicative of multi-band or edge artifacts)
2. Time series showing

a. Sudden, abrupt changes
3. Power spectrum

a. Main frequency above 0.1 Hz
4. Multi-echo information

a. High rho or low kappa

When classification based on these criteria was unclear, the component was accepted.

#### 2.2.4. General linear models for denoising

The SE and ME-OC images from each functional dataset were processed using AFNI’s 3dREMLfit, applied to the signal from each voxel *(Y_SE_* for single-echo data and *Y_OC_* for multi-echo optimally combined data).

1. The *SE model* included six motion parameters from volume realignment *(Mot)*, up to fourth-order Legendre polynomials *(Poly)*, the end-tidal CO_2_ regressor (*CO_2_),* and left- and right-hand grip regressors *(RGrip* and *LGrip)*.

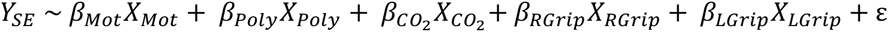
2. The *ME-OC model* included 6 motion parameters from volume realignment *(Mot)*, up to fourth-order Legendre polynomials *(Poly)*, the end-tidal CO_2_ regressor (*CO_2_),* and left- and right-hand grip regressors *(RGrip* and *LGrip)*.

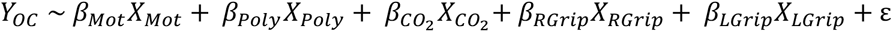
3. The *ME-ICA model* included 6 motion parameters from volume realignment, up to fourth-order Legendre polynomials, the end-tidal CO_2_ regressor (*CO_2_),* left- and right-hand grip regressors *(RGrip* and *LGrip)*, and the rejected ME-ICA components *(IC_rej_)*.

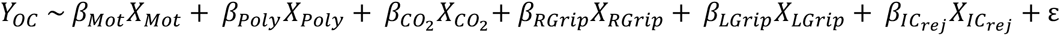

For each of the three models, a denoised dataset was also calculated by subtracting the fitted timeseries associated with the nuisance regressors (i.e., the product of the corresponding beta-coefficient maps and the regressor time series) from the Y_SE_ and Y_OC_ data. Nuisance regressors are defined here as all the model regressors other than *RGrip* and *LGrip*.

#### 2.2.5. Quantification of head motion

Framewise displacement (FD) was calculated from the volume realignment parameters as the sum of the absolute difference of the realignment parameters between samples (Power et al., 2012). The motion parameters are defined as X (anterior-posterior axis), Y (left-right axis), Z (inferior-superior axis), Roll (rotation around X), Pitch (rotation around Y), and Yaw (rotation around Z). Translation motion was calculated in millimeters; rotation motion was calculated in degrees and converted to millimeters before using to calculate FD, as done previously (Power et al., 2012). The degrees to millimeters conversion was performed by calculating displacement on the surface of a sphere; 50 mm was used as the sphere radius since it is the approximate distance from the cerebral cortex to the center of the head. As described in *2.2.3*, for SE data, these volume realignment parameters were calculated with respect to the second echo Single Band reference image; for ME data, these volume realignment parameters were calculated with respect to the first echo Single Band reference image. To see how many data points may have been removed if censoring was applied, the number of data points with a FD > 0.5 was calculated. While several censoring thresholds have been used previously, FD > 0.5 is within the range of previous fMRI studies (Power et al., 2014, 2017; Siegel et al., 2014), and is used here as an example. The left- and right-hand grasp force traces were summed to create an overall task timeseries. Pearson correlation coefficients comparing this task timeseries with each of the six motion parameters from volume realignment were calculated.

#### 2.2.6. Assessment of head motion dissociation from Blood Oxygenation Level Dependent (BOLD) signal and noise reduction

As done by Moia et al. (2021), the denoised datasets described in *2.2.4* were used to calculate DVARS, the spatial root mean square of the first derivative of the signal (Smyser et al., 2010) using 3dTto1D (AFNI). For SE, ME-OC, and ME-ICA models, DVARS was calculated within the tighter brain extraction mask created after tedana. A dataset with high association between head motion and BOLD signals would display greater DVARS with increasing FD, while a dataset with low association between head motion and BOLD signals would display consistent DVARS with increasing FD. To quantify this association, the slopes of the DVARS vs. FD relationship were calculated for each scan (DVARS ∼ FD). For the Limited and Amplified motion conditions, the slopes of the DVARS vs. FD relationship were modeled at the group level as Slope ∼ Model + (1|Subject).

The denoised datasets described in *2.2.4* were also used to create gray plots to visually assess the level of noise present in the scans (Power, 2017). The anatomical image for four representative datasets was segmented into gray matter and white matter (fsl_anat) and thresholded at 0.5. The gray and white matter segmentations were then linearly transformed to functional space (epi_reg, FLIRT, FSL). 3dGrayplot (AFNI) was used to create gray plots for each scan within the gray matter and white matter segmentations, and voxels were ordered by how well they matched the two leading principal components within each tissue type.

#### 2.2.7. Spatial correlation analysis

Voxel-wise spatial correlation analysis was performed in R (version 2022.07.1) to compare the hand grasp activation maps from the Limited and Amplified motion scans within each subject. The Limited motion scan served as an estimate of the ground-truth activation for a given subject. Beta coefficient maps corresponding to the *RGrip* and *LGrip* regressors were converted to MNI space by applying a single concatenated spatial transformation of the functional images to the subject’s T1-w structural image (epi_reg, FSL) and from this image to the FSL 2-mm MNI template (FLIRT and FNIRT, FSL). Transformed maps were masked with the 2-mm MNI brain mask provided by FSL. The Pearson spatial correlation coefficients (r) between the Limited and Amplified activation maps were calculated for each subject and hand, then converted to Fisher z values.

The Limited motion activation maps were thresholded to find areas of positive activation at p_FDR_ < 0.05 to highlight significant areas of activation in the ‘ground-truth’ dataset. Similarly, the Pearson spatial correlation coefficient (r) between the Limited and Amplified scans was calculated within a mask of these areas of significant positive activation, then converted to Fisher z values. Significant differences in the spatial correlations between SE, ME-OC, and ME-ICA models were tested as FisherZ ∼ Model + (1|Subject).

To better understand agreement of areas of activation between Limited and Amplified maps, the Dice similarity coefficient was also calculated between regions of significant positive activation for the two motion conditions. Both Limited motion and Amplified maps were thresholded at p_FDR_ < 0.05. Significant differences in the Dice coefficients between SE, ME-OC, and ME-ICA models were tested as Dice ∼ Model + (1|Subject).

#### 2.2.8. Subject-level ROI analysis

To facilitate quantitative comparisons of motor activation across subjects and scans, several ROIs known to be involved with hand grasping force generation were identified using manual and automated methods.

The right- and left-hand knob areas of the motor cortex, corresponding to hand motor activation (Yousry et al., 1997), were manually drawn for each participant in anatomical space, then non-gray matter voxels removed by finding overlap with a gray matter mask created using fsl_anat (thresholded at 0.5). This mask was dilated by two voxels, then linearly transformed to the functional space for each subject (epi_reg, FLIRT, FSL).

Right and left motor cerebellum ROIs were created from the SUIT probabilistic cerebellum atlas in MNI space (J. Diedrichsen et al., 2011; Jörn Diedrichsen et al., 2009). Hand and finger motor tasks have been associated with activity in ipsilateral cerebellum lobules V/VI and VIIIa/b (Ashida et al., 2019; Spencer et al., 2007; Stoodley, 2012). Consequently, right lobules V/VI and VIIIa/b from the SUIT atlas were thresholded at 50%, binarized, and combined to form a right motor cerebellum ROI; the same was done with the left motor lobules to create a left motor cerebellum ROI. These masks in MNI space were then non-linearly transformed to functional space for each subject (FNIRT, FSL).

The percent positive activated voxels (defined as voxels showing a statistical significance of p_FDR_ < 0.05), median positive beta coefficient, and median positive t-statistic corresponding to the *RGrip* regressor were calculated in the left-hand knob and right motor cerebellum ROIs for each scan in functional space. The percent positive activated voxels (p_FDR_ < 0.05), median positive beta coefficient, and median positive t-statistic corresponding to the *LGrip* regressor were calculated in the right-hand knob and left motor cerebellum ROIs.

Significant differences in the parameters calculated by each model were modeled as Parameter ∼ Model + (1|Subject). Significant differences in the median beta coefficient calculated for the Limited and Amplified conditions were modeled as BetaCoefficient ∼ Condition + (1|Subject). The relationships of the median beta coefficient with FD and with task correlation in the X direction (Xcorrelation) were modeled as BetaCoefficient ∼ FD and BetaCoefficient ∼ Xcorrelation, respectively.

#### 2.2.9. Correlation of motor task regressors with end-tidal CO_2_ regressor

As subjects might modify their breathing during the hand grasps, which in turn alters the level of arterial CO_2_ and thus influences blood flow and BOLD fMRI signals throughout the brain due to vasodilation (Birn et al., 2006; Farthing et al., 2007), we accounted for this potential confounding effect by including an end-tidal CO_2_ regressor in our subject-level models (*2.2.4*). End-tidal CO_2_ recordings are a reasonable non-invasive estimate of arterial CO_2_ levels (McSwain et al., 2010). We assessed the task-correlation of respiratory effects in our datasets by calculating Pearson correlation coefficients comparing the left- and right-hand grip regressors and the sum of these grip regressors with the end-tidal CO_2_ regressor. To better understand the effect of adding this CO_2_ regressor to our models, we analyzed the datasets with additional subject-level general linear models that did not include the CO_2_ regressor.

#### 2.2.10. Motor activation group analysis

Group-level activation maps were calculated for the Limited and Amplified motion scans to compare motor activation results across subjects with low and high amounts of motion, respectively.

Beta coefficient and t-statistic maps for *RGrip* and *LGrip* for each subject were converted to MNI space by applying a single concatenated spatial transformation of the functional images to the subject’s T1-w structural image (epi_reg, FSL) and from this image to the FSL 2-mm MNI template (FLIRT and FNIRT, FSL). The quality of these registration steps is shown in **Supplemental Figure 2**. Group-level analysis was performed using AFNI’s 3dMEMA (Chen et al., 2012), using contrasts for *RGrip* > 0 and *LGrip* > 0. Group-level maps were thresholded at p < 0.005 and clustered at α < 0.05, using a bi-sided t-test (3dFWHMx, 3dClustSim, 3dClusterize, AFNI).

A paired group-level analysis was conducted between the Limited and Amplified motion conditions for each of the three models (3dttest++, AFNI). This analysis was conducted on the *RGrip* and *LGrip* activation maps separately.

Hand-knob and cerebellum motor area ROIs used for subject-level quantitative analysis (*2.2.8*) were similarly created for group-level ROI analysis in MNI space. The right- and left-hand knob areas of the motor cortex were manually drawn in MNI space, non-gray matter voxels removed by finding overlap with a gray matter mask, then dilated by two voxels. Motor cerebellum areas were created from the SUIT atlas as described in *2.2.8*.

In each ROI, the percent positive activated voxels were calculated as those in the significant clusters found as described above, and the median positive beta coefficient and median positive t-statistic were calculated using the unthresholded group-level activation maps.

### 2.3. Stroke datasets

Our head motion simulation for the Amplified condition aimed to approximate the head motion characteristics of motor-impaired participants. Thus, we analyzed head motion data from two participants with chronic hemiparetic stroke and unilateral hand impairment to compare with our healthy participant simulation results. Stroke Subject 1 (M, 65yrs) experienced impairment of the left hand, and Stroke Subject 2 (M, 65yrs) experienced impairment of the right hand. Two fMRI datasets were collected from each participant and used only to measure head-motion realignment parameters. No further analysis was performed on these datasets, as it was outside the scope of the current study. Detailed scan parameters for the stroke participants can be found in **Supplemental Table 1**.

The stroke participants performed the same motor task described above and were instructed to keep their head as still as possible (note, the first participant performed 7 left hand grips and 9 right hand grips). Head-motion realignment parameters were computed with reference to the Single Band reference images taken at the start of each scan. (3dvolreg, AFNI).

## 3. Results

All structural and functional scans were successfully collected as described above. The Amplified condition scan from Subject 1 was excluded from spatial correlation, ROI, and group activation analysis because the average head motion (FD) of the scan was identified as an outlier (**Supplemental Figure 3**). Results that include the Amplified condition scan from Subject 1 can be found in Supplemental Materials.

### 3.1. Motion characteristics of the datasets

The forces produced by all healthy and stroke subjects are shown in **Figure 1A** and **Figure 1B**. All healthy participants had high task accuracy, demonstrated by force traces thatstayed within the green shaded area representing the target force range (35-45% MVC). The stroke participants were also able to successfully perform the task with both paretic and non-paretic hands. Stroke Subject 2 demonstrated some concurrent grasping with the hand that was not instructed by the task, and also missed one grasping period during the first scan. A non-zero baseline force, seen in some of the subjects’ force traces, may have been due to the participant lightly grasping the load cell during the rest periods.

**Figure 1.**
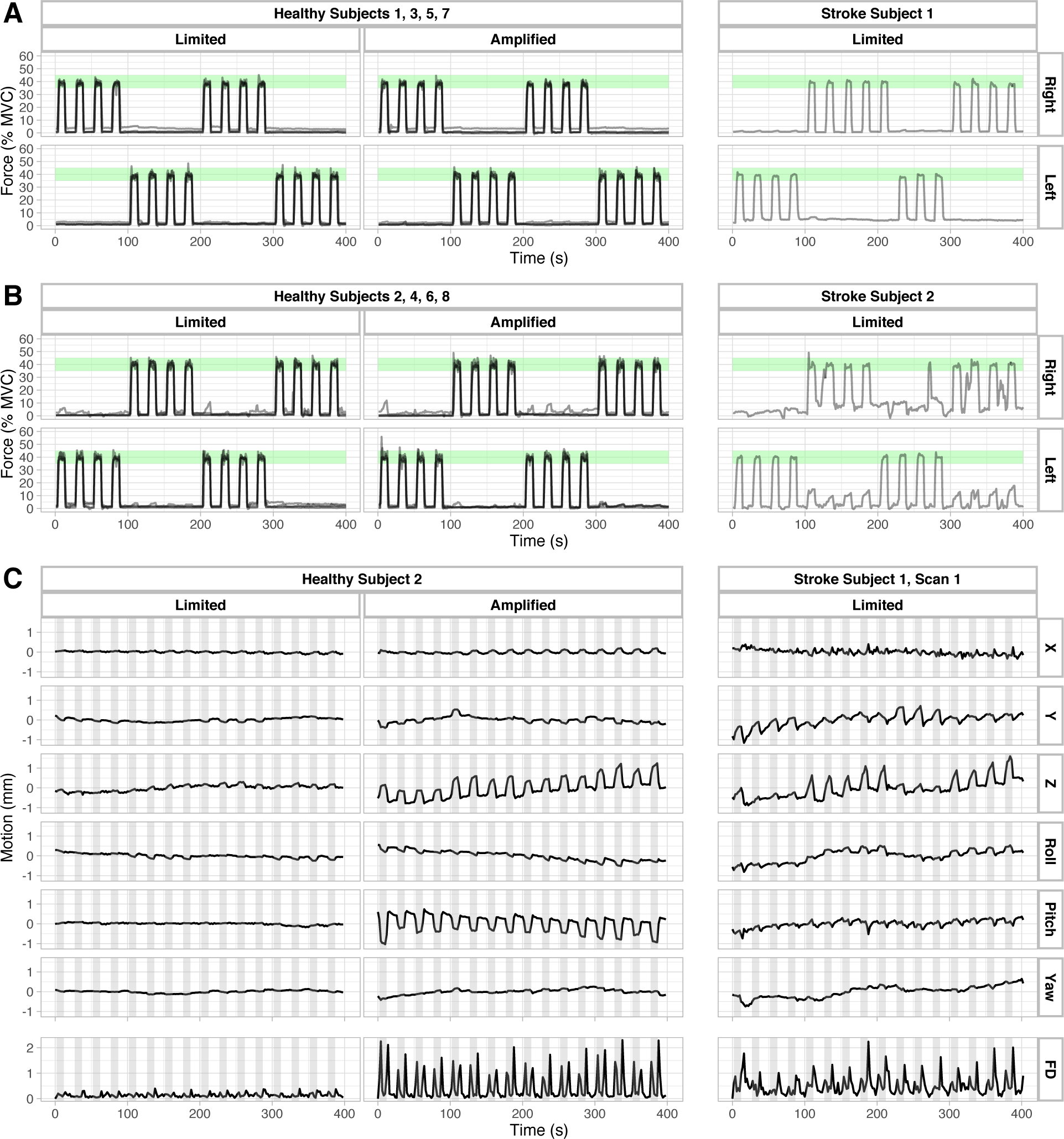
(A) Force traces from healthy subjects 1, 3, 5, and 7 and Stroke Subject 1, normalized to each subject’s hand grasp at maximum voluntary contraction (MVC). Green shaded areas indicate target force range (35-45% MVC) displayed in visual task instructions. (B) Force traces from healthy subjects 2, 4, 6, and 8 and Stroke Subject 2, normalized to each subject’s MVCs. Green shaded areas indicate target force range (35-45% MVC) displayed in visual task instructions. (C) Motion parameters from volume realignment and calculated Framewise Displacement (FD) from one representative healthy subject (Subject 2) in the Limited motion and Amplified motion conditions and one Stroke subject (Stroke Subject 1, Scan 1) with mild motor impairment. Gray shaded areas represent periods when the subjects were instructed to perform the grasping task.

**Figure 1C** displays the motion parameters from volume realignment and calculated FD for one representative healthy subject (Subject 2) and one stroke subject (Stroke Subject 1, Scan 1) during the hand grasp task. Head motion was observed to be highly related to the motor task timings, which are indicated by the gray shaded regions. This task-correlation effect is visible even in the Healthy Limited condition, though with lower amplitudes of motion. During the Amplified condition, the healthy subject demonstrated increased task-correlated head motion in the Z and pitch directions. In contrast, the stroke subject primarily displayed task-correlated motion in the Y and Z directions.

These observations are quantified in **Figure 2**, which summarizes the average FD, percent of volumes that would be censored (FD > 0.5), and the correlation between the estimated realignment parameters and hand grasping task for each scan. Every healthy participant had an increase in FD and the percentage of potentially censored volumes during their Amplified motion scan compared to the Limited motion scan. Generally, the healthy participants showed an increase in task-correlated head motion in the Amplified motion scan, particularly in the X, Z, and Pitch directions that are associated with the added, self-induced head nodding. Note that participants with a high increase in FD between scans did not necessarily have a high increase in task-correlated head motion, and vice versa, demonstrating the important distinction between these two characteristics of head motion during fMRI (ex. Subject 1 Amplified, Subject 4 Amplified). The two scans for Stroke Subject 1 have similar average head motion and percent potentially censored volumes to that of the healthy Amplified scans, while Stroke Subject 2 had much higher average head motion. Both Stroke Subjects 1 and 2 displayed task-correlated head motion, though the correlations were somewhat more similar to the healthy Limited scans. FD calculations across the scans showed high similarity between the SE and ME analyses (**Supplemental Figure 4**).

**Figure 2.**
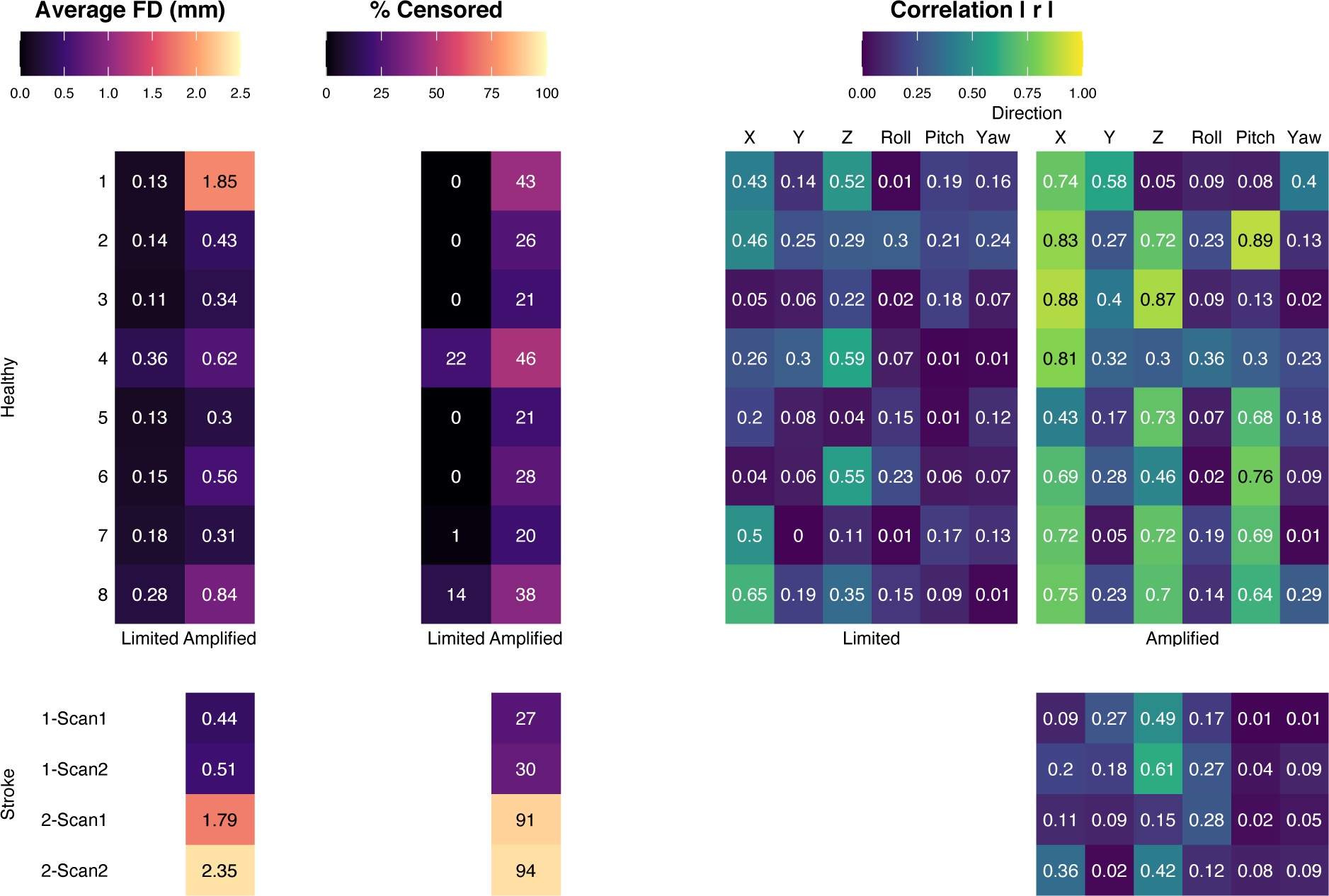
Average Framewise Displacement (FD), percent of data points censored at FD > 0.5, and correlation of the task regressor (LGrip + RGrip) with the timeseries of the realignment parameters for each functional scan. For healthy participants, the Limited and Amplified motion tasks are shown. For stroke participants, the two scans acquired in each participant are shown, during which the instructions were to keep head motion minimal.

### 3.2. Assessment of head motion dissociation from Blood Oxygenation Level Dependent (BOLD) signal and noise reduction

The ability of each model to dissociate the effects of head motion from the BOLD fMRI signal was assessed by examining the relationship between DVARS, a measure of instantaneous fMRI signal change, and FD, a measure of head motion, at each fMRI timepoint. An ideal denoising strategy would minimize the magnitude of this relationship, meaning large changes in BOLD signal would not occur at the same times as large changes in head position. **Figure 3A** shows DVARS vs. FD plots for each scan, calculated for all three models, in the Limited and Amplified conditions. The ME-ICA models consistently had lower DVARS than the SE and ME-OC models, even during volumes with high FD. **Figure 3B** summarizes the DVARS vs. FD relationship across scans. Within both the Limited and Amplified motion conditions, the ME-ICA model had significantly lower slopes than the SE and ME-OC models (p < 0.05, Bonferroni-corrected), indicating a better ability to dissociate the effects of head motion from the BOLD signal. Of note, when the absolute value of the slopes is compared, there is no longer a significant difference between the ME-OC and ME-ICA models in the Limited motion condition.

**Figure 3.**
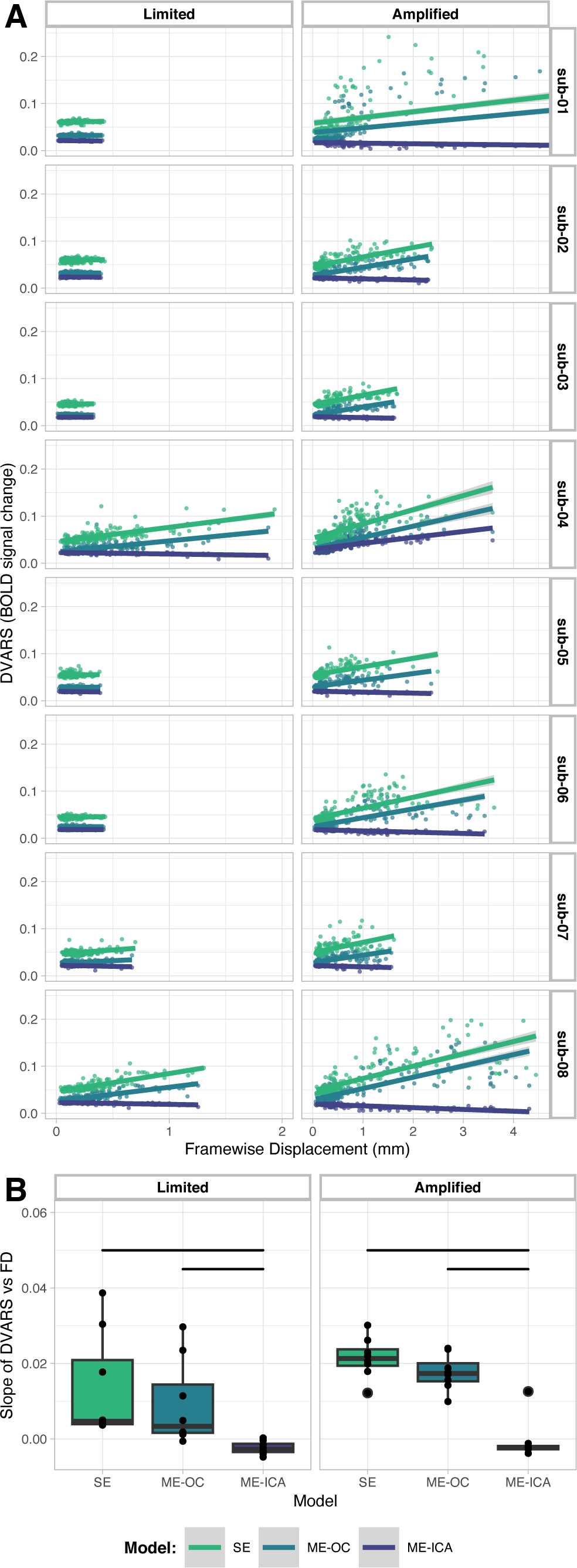
(A) Relationship between DVARS and Framewise Displacement (FD) for each scan. A more horizontal fit line indicates lower correlation of the change in BOLD signal with the amount of head motion at each time point of the scan. Note difference in X-axis scale between Limited and Amplified conditions. (B) Summary of slopes from DVARS vs. FD graphs in (A). Bars indicate significant difference between models (p < 0.05, Bonferroni-corrected).

The effect of each model on the noise present in the voxels was also visually assessed by examining gray plots of representative scans across the range of motion. An effective denoising model would reduce noise in regions not associated with the task (white matter) and retain BOLD signal changes in regions associated with the task (gray matter). **Figure 4** shows gray plots for four representative scans with low, moderate, high, and very high levels of head motion. The timeseries shown were denoised by the SE, ME-OC, and ME-ICA models. At low and moderate levels of head motion, the three models show a similar ability to retain BOLD signal changes in gray matter while minimizing signal changes in white matter. In the high and very high motion scans, the SE and ME-OC models display widespread signal changes in gray and white matter that are time-locked with large changes in FD. The ME-ICA model greatly reduces these motion-related changes in white matter and retains BOLD signal changes in gray matter induced by the task-related activity.

**Figure 4.**
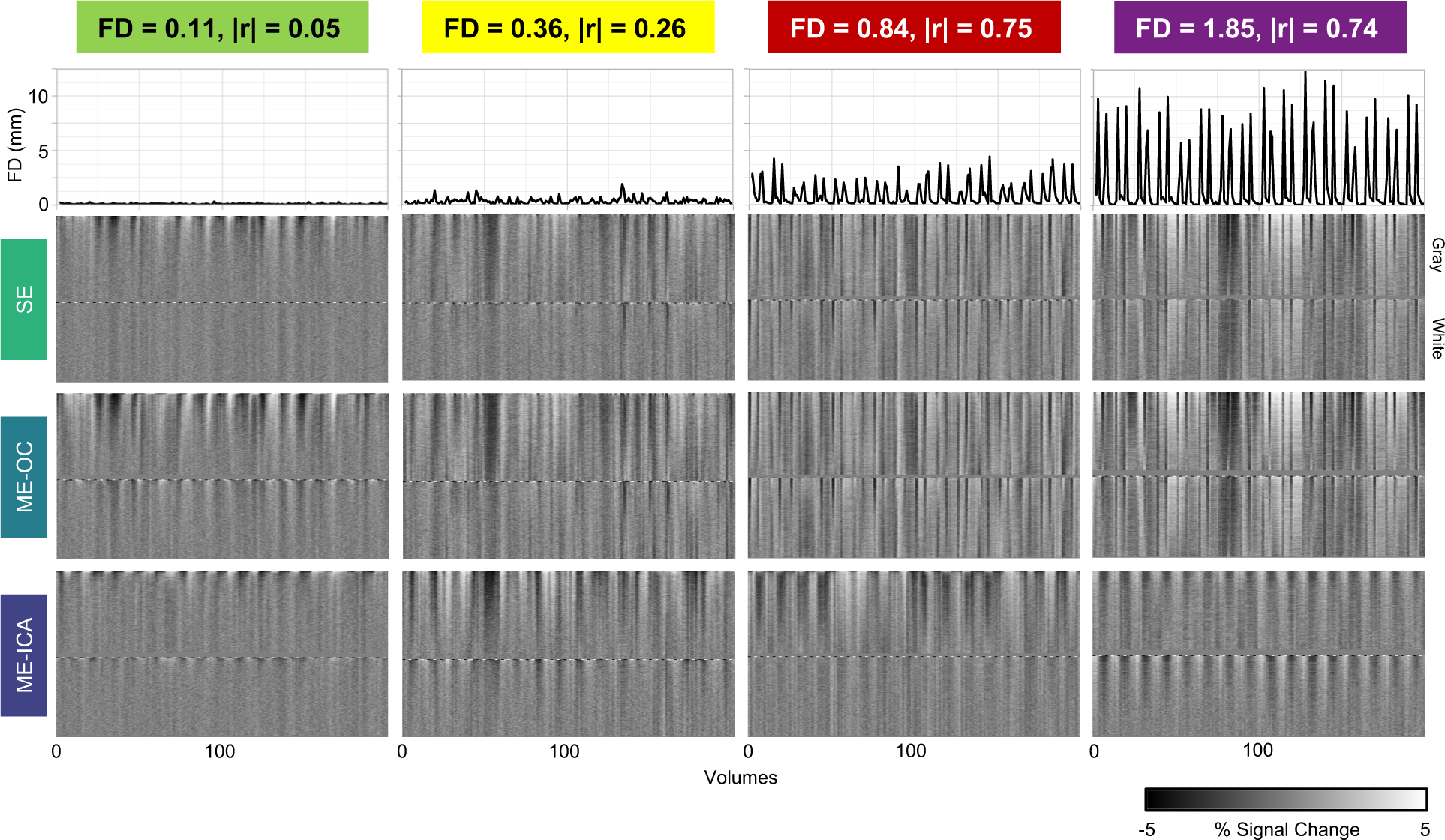
Gray plots and Framewise Displacement (FD) across the scans, using single-echo, multi-echo, and multi-echo ICA denoised timeseries. Four representative scans with low, moderate, high, and very high levels of head motion are shown (Subject 3 Limited, Subject 4 Limited, Subject 8 Amplified, and Subject 1 Amplified, respectively). Average FD and correlation of task with motion in the X direction for each scan are listed. FD was calculated using volume realignment parameters obtained from the first echo. Gray plots were calculated with AFNI’s 3dGrayplot. The signals shown were from voxels within the gray matter and white matter regions, and voxels are ordered by how well they match the two leading principal components within each tissue type. ME-ICA denoising reduced ordered noise related to head motion. Specifically, in higher motion datasets, ME-ICA can be seen to reduce noise in white matter regions, and retain BOLD changes related to the task within the gray matter.

### 3.3. Subject-level activation

We analyzed subject-level activation maps to understand how task-correlated head motion affects our ability to visualize motor activation and evaluate how ME preprocessing (ME-OC and ME-ICA) can reduce these confounding effects. **Figure 5** shows subject-level activation maps for the right grip task, with location of slices chosen to depict activation in the hand motor area. Four representative scans with low, moderate, high, and very high levels of average head motion (average FD) are depicted. In the low motion scan, typical of healthy participants in the Limited motion dataset, there is minimal difference between the three models; activation in the motor cortex is similar across models and noise is slightly reduced in the ME models, seen as apparent smoothing across the brain and reduced artifacts at the edges of the brain and cerebellum. In the moderate motion scan, the SE model shows noise at the edges of the brain, which is mitigated using the ME models. In the high motion scan, the SE model shows both noise at the edges of the brain and a banding artifact that may be related to the interaction between the multi-band slice acquisition and head motion. The ME-OC model does not remove these artifacts, while the ME-ICA model is able to better mitigate them. This same effect is observed to a greater degree in the very high motion scan, which was an outlier for the healthy Amplified datasets in terms of the average FD, though comparable to the motion observed in Stroke Subject 2. Activation in the motor cortex, indicated by arrows in **Figure 5**, is present in all scans and models; however, in the high and very high motion scans, motor cortex activation is most easily visually differentiated from the surrounding region when using ME-ICA. The moderate, high, and very high head motion scans are more similar to the motion of the stroke participant scans, and ME-ICA may have particular utility in reducing the larger motion-related artifacts in these scans. Similar results were observed for the rest of the scans.

**Figure 5.**
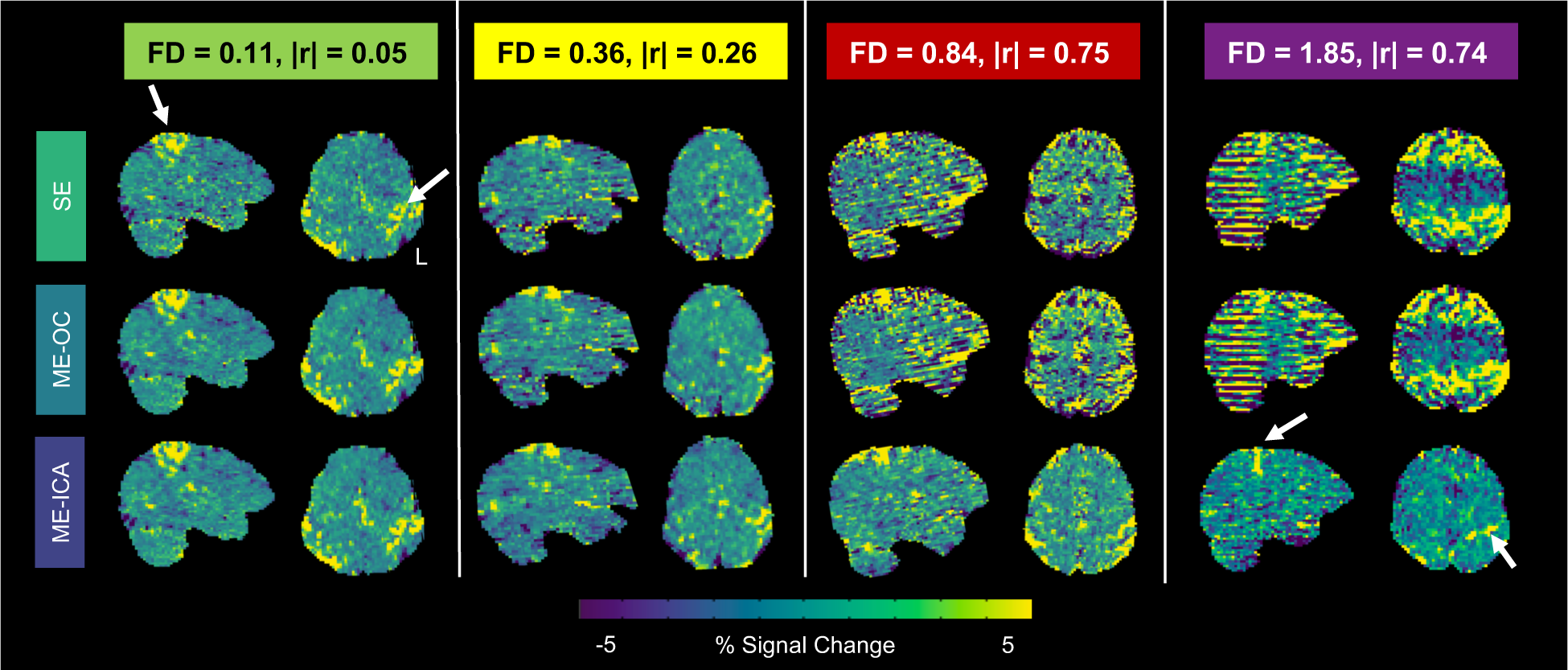
Subject-level maps in functional space of percent BOLD signal change related to the right grip task. Four representative scans with low, moderate, high, and very high levels of head motion are shown (Subject 3 Limited, Subject 4 Limited, Subject 8 Amplified, and Subject 1 Amplified respectively). Average FD and correlation of task with motion in the X direction for each scan are listed. Arrows indicate expected regions of activation in the motor cortex. As head motion increases, artifacts increase at the edges of the brain and banding patterns appear. These artifacts are mitigated by ME-OC and ME-ICA.

### 3.4. Spatial correlation between Limited and Amplified motion activation maps

To quantify the similarity of hand grasp activation estimates across different levels of motion, we conducted a within-subject spatial correlation analysis. The Limited motion scan served as an approximate ground-truth activation map for each subject. A higher similarity of Limited and Amplified motion activation maps within a subject indicates a better ability of the model to provide stable activation estimates when motion-related artifacts are increased. **Figure 6A** shows subject-level activation maps related to the right grip task performed with Limited and Amplified motion, across three representative scans with high, moderate, and low levels of spatial correlation between Limited and Amplified maps. The spatial correlation improves with ME-OC compared to SE, and with ME-ICA compared to ME-OC. The spatial correlation patterns across all subjects are shown in **Figure 6B**. When unthresholded activation maps are compared, the ME models lead to significantly higher spatial correlations than SE. Notably, the RGrip activation map from Subject 7 leads to a negative spatial correlation with ME-ICA. For Subject 7, all models show regions of negative activation in the Limited motion map that are positive in the Amplified motion map. This effect is most prominent with the ME-ICA model, leading to a negative spatial correlation. However, the relevant regions are outside motor areas and have relatively low t-statistics that do not pass thresholding at p_FDR_ < 0.05. The activation maps from Subject 7 are shown in **Supplemental Figure 5**.

**Figure 6.**
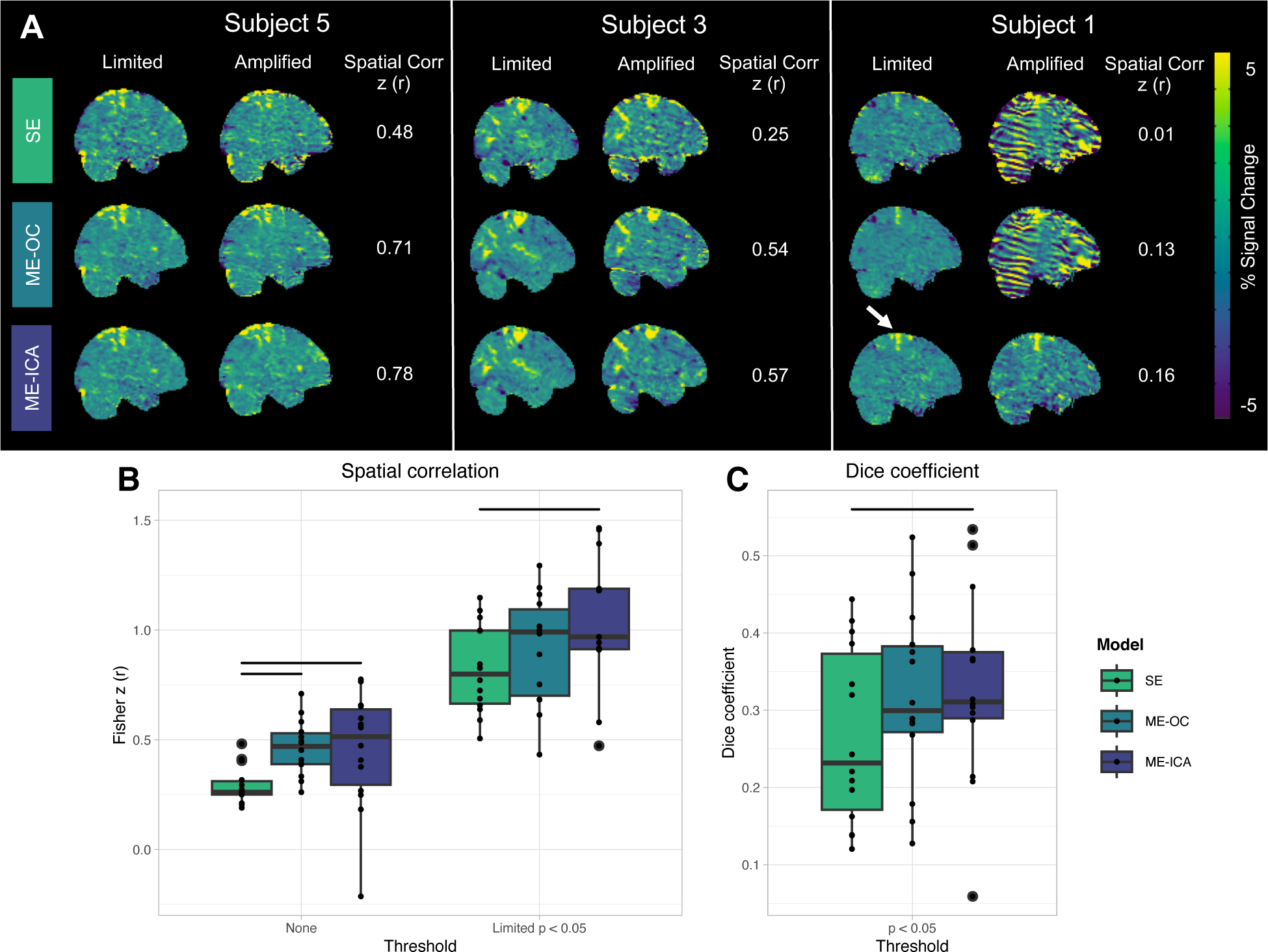
(A) Subject-level maps in MNI space of percent BOLD signal change related to the right grip task performed with Limited and Amplified motion. Three representative scans with high, moderate, and low levels of spatial correlation between Limited and Amplified maps are shown. Pearson r correlation values shown were calculated between unthresholded maps, then converted to Fisher z. The arrow indicates the expected region of activation in the motor cortex. (B) Spatial correlation between Limited and Amplified maps. Correlation was calculated between unthresholded maps and between maps that were thresholded by voxels with significant positive activation in the Limited map. Significant positive activation was determined by thresholding the Limited map at p_FDR_ < 0.05, using a t-statistic accounting for degrees of freedom in the model. Only areas of positive activation were retained. A mask of these significant positive voxels was applied to Limited and Amplified beta parameter maps before correlation was calculated. Maps were thresholded to show correlation after reducing effects of background noise voxels on spatial correlation estimates. Subject-level maps in MNI space from RightGrip and LeftGrip activation were included. (C) Dice similarity coefficients calculated between areas of significant positive activation in Limited and Amplified scans, respectively. Significant voxels were found as in (B), independently calculated for Limited and Amplified scans. Subject-level maps in MNI space from RightGrip and LeftGrip activation were included. Dots represent values from individual scans. Bars indicate significant difference between models (p < 0.05, Bonferroni-corrected). Results that include motion outlier Subject 1 in group-level visualization and identify individual subjects by color can be found in **Supplemental Figure 6**.

To focus on areas of activation and reduce effects of background noise, such as in the case of Subject 7, the Limited motion activation maps were thresholded to find areas of significant positive activation at p_FDR_ < 0.05. Spatial correlation was then calculated between Limited and Amplified maps within these regions of significant activation (**Figure 6B**). When focusing on these regions, ME-ICA still led to a significantly higher spatial correlation than the SE model. This result suggests that, when ME-ICA is implemented, there is higher within-subject agreement between the activation calculated from a high motion scan and the ‘ground-truth’ activation from the Limited motion scan.

A comparison was also completed between the areas of significant positive activation in the Limited motion maps and the areas of significant positive activation in the Amplified motion maps. A Dice similarity coefficient was found between these regions for each subject and hand. Again, the ME-ICA model led to a significantly higher Dice coefficient than the SE model. Across all models, the Dice coefficient values were low, with most values below 0.5. A key reason for these low values is that the voxel-wise thresholding performed before comparison leaves scattered activated voxels outside the expected motor regions for each scan. These voxels vary between Limited and Amplified motion scans, lowering the overall Dice coefficient despite agreement in motor regions. However, the overall trend seen across models is still thought to represent better agreement of maps when using ME-ICA compared to SE, as the trend is also comparable to the observations seen through the aforementioned spatial correlation analyses.

### 3.5. Subject-level ROI analysis

To quantify the differences in motor task activation across scans and analyses, activation characteristics were specifically probed in brain regions related to hand motor tasks: the hand-knob regions of the motor cortex and cerebellum lobules V/VI and VIIIa/b. The manually drawn hand-knob masks for each subject are shown in **Supplemental Figure 7**. For each motor task (left- or right-hand targeted grasping), activation results were analyzed in the contralateral hand knob and ipsilateral cerebellum. **Figure 7A** shows the subject-level activation extent, median positive beta coefficient, and median positive t-statistic in each region, summarized across the Limited and Amplified motion conditions and the three models. Activation extent was defined as the percent positive activated voxels (p_FDR_ < 0.05) in each ROI.

**Figure 7.**
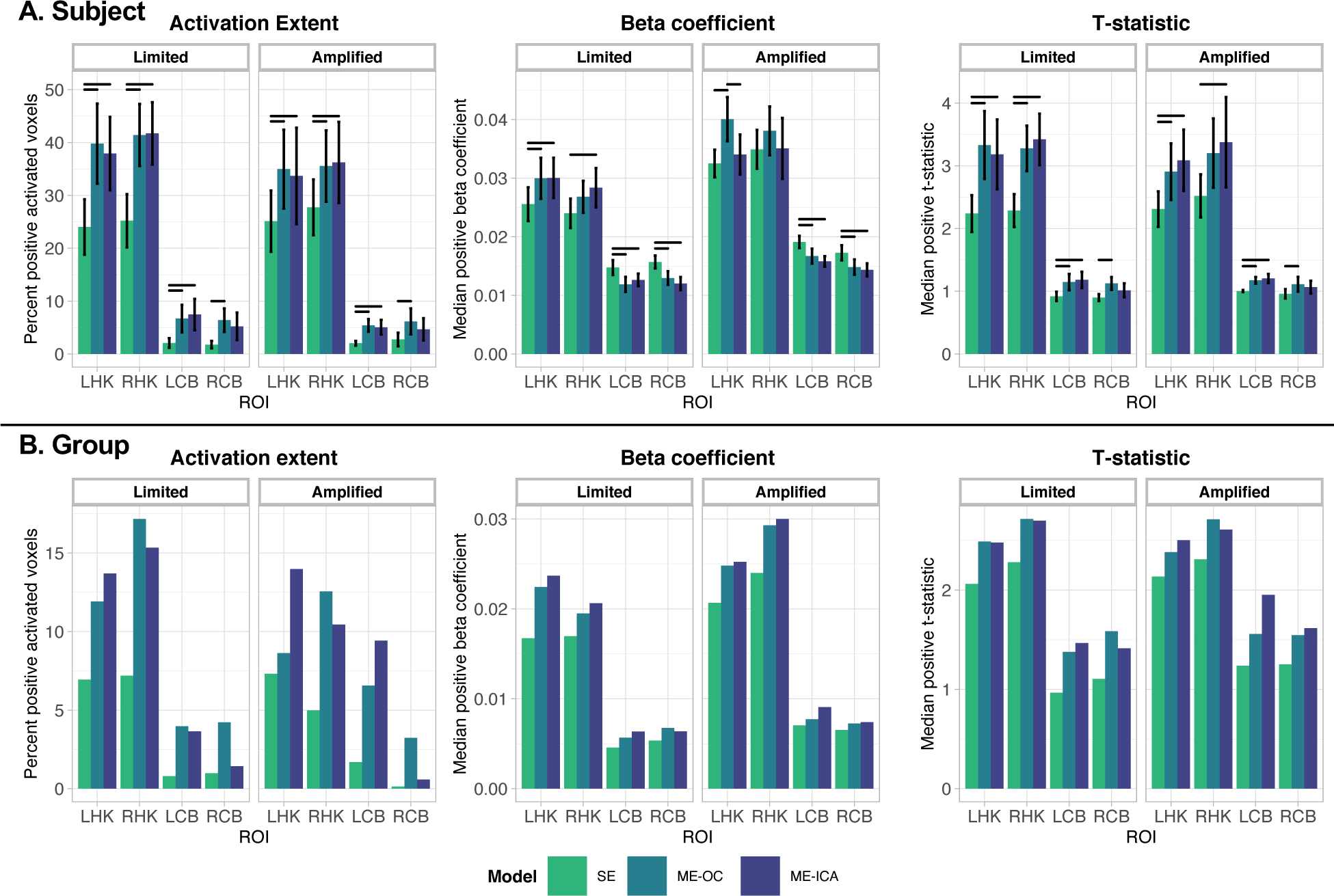
(A) Subject-level activation extent, beta coefficient, and t-statistic values for each scan associated with response to hand grasp tasks. Vertical bars indicate standard error across subjects. Horizontal bars indicate significant difference between models (p < 0.05, Bonferroni-corrected). (B) Group-level activation extent using clustered group activation maps, and beta coefficient and t-statistic values for unthresholded group activation maps associated with response to hand grasp tasks. Group maps were calculated with 3dMEMA, AFNI. In (A) and (B), values are shown for four ROIs: values associated with right hand grip in the left hand knob (LHK) and right cerebellum hand motor areas (RCB) and values associated with left hand grip in the right hand knob (RHK) and left cerebellum hand motor areas (LCB). Results that include the Subject 1 Amplified scan in analysis and show individual subject data points can be found in Supplemental Figure 8.

In the hand-knob regions, the ME models had significantly greater activation extent than SE (p < 0.05). In the cerebellum, both ME models had greater activation extent than SE, with only the ME-OC achieving significance in the right cerebellum. These results are seen in both the Limited and Amplified motion conditions.

In terms of beta coefficients, with Limited motion, the ME models had greater median positive beta coefficients than SE in the hand-knob regions, but lower beta coefficients than SE in the cerebellum. With Amplified motion, ME-OC leads to a higher beta coefficient than both SE and ME-ICA; this trend is significant in the left hand knob, though also present in the right hand knob. Given that the hand grasp task was the same across the Limited and Amplified motion scans, the true activation represented by the beta coefficient within the hand-knob region is expected to be similar. In the left hand knob, the median beta coefficient is significantly different between the Amplified and Limited conditions for only the ME-OC model (SE: p = 0.07; ME-OC: p = 0.03; ME-ICA: p = 0.38). In the right hand knob, the median beta coefficient is significantly different between the Amplified and Limited conditions for SE and ME-OC (SE: p = 0.03; ME-OC: p = 0.04; ME-ICA: p = 0.16). For the Amplified motion scans, the ME-OC model’s higher beta coefficients suggest a possible inflation of activation effect size using ME-OC, mitigated by the addition of ME-ICA. The Amplified to Limited condition comparisons are more mixed for left cerebellum (SE: p = 0.002; ME-OC: p < 0.0001; ME-ICA: p = 0.02) and right cerebellum (SE: p = 0.13; ME-OC: p = 0.16; ME-ICA: p = 0.01), with no clear pattern across models.

The median positive t-statistic was larger in the ME conditions compared to SE (**Figure 7A**). During Amplified motion scans, ME-ICA had comparable t-statistics to ME-OC in the hand-knob regions, though the beta coefficient was smaller. Therefore, the comparable t-statistic may reflect a large decrease in the variance of the residuals in the ME-ICA model compared to ME-OC, suggesting a better ability of the ME-ICA model to mitigate uncertainty in the model.

We also plotted the relationship of the beta coefficient in each ROI with FD and with task correlation with motion in the X direction (**Figure 8**). A slope closer to zero indicates a lower relationship of the beta coefficient with these motion-related metrics. This analysis was performed across all Limited and Amplified scans, excluding the motion outlier Subject 1, to investigate the impact of ME-ICA across the range of motion available in this dataset. For all but one case (right cerebellum, X correlation), ME-ICA resulted in the lowest relationship of the beta coefficient with average motion and task-correlation of motion, compared to SE and ME-OC. This finding suggests that ME-ICA may perform better at maintaining the consistency of the beta coefficient across a dataset, independent of the degree of head motion in a given scan.

**Figure 8.**
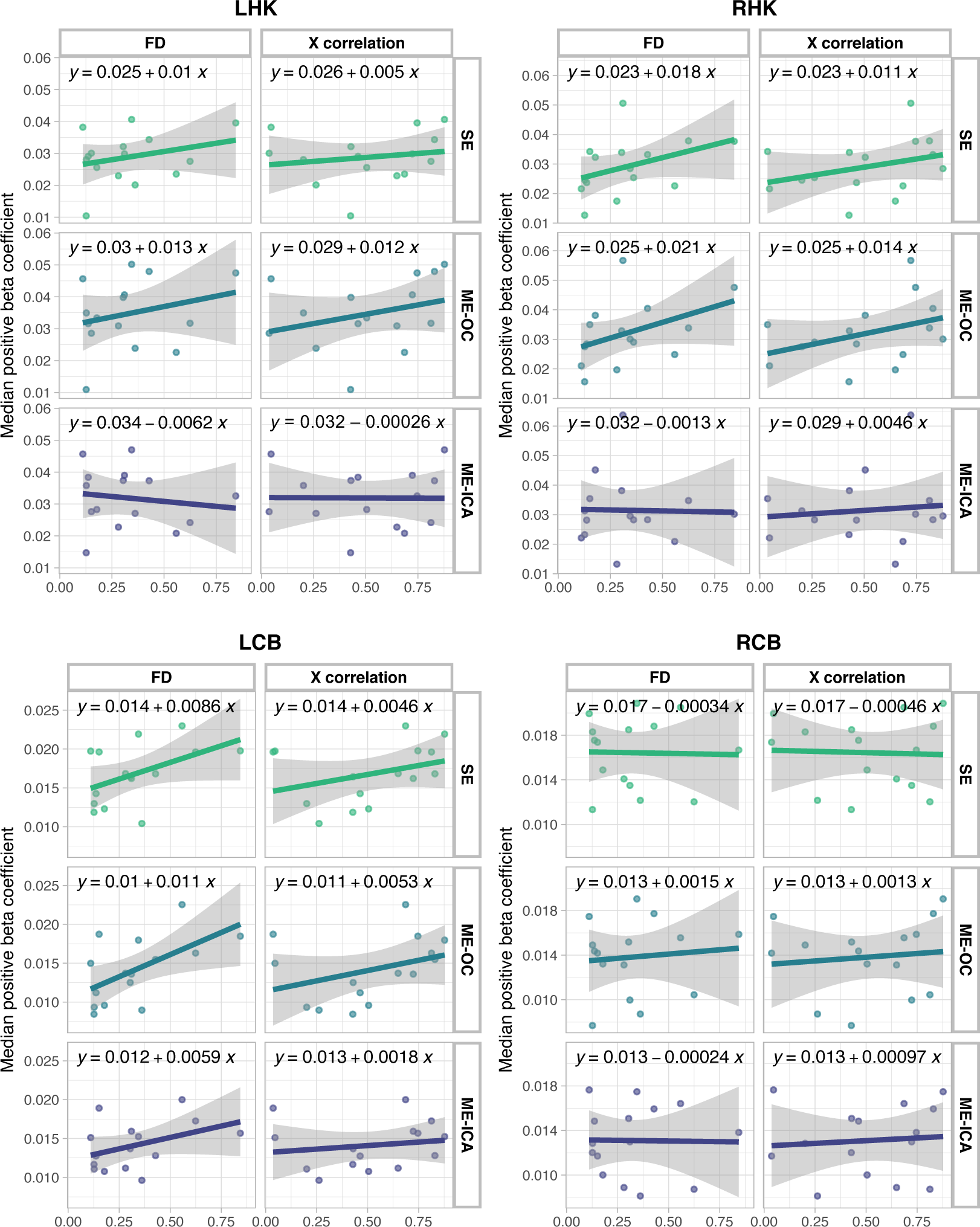
Median beta coefficients calculated using SE, ME-OC, and ME-ICA models shown for each Limited and Amplified motion scan, plotted against the average FD (mm) and against the task correlation with motion in the X direction for each scan. Beta coefficient values are shown for four ROIs: values associated with right-hand grip in the left hand knob (LHK) and right cerebellum hand motor areas (RCB) and values associated with left-hand grip in the right hand knob (RHK) and left cerebellum hand motor areas (LCB). For all but one case (RCB, X correlation), ME-ICA results in the lowest relationship (slope) of the beta coefficient with average motion and task-correlation of motion. Results that include the Subject 1 Amplified scan in analysis can be found in **Supplemental Figure 9**.

### 3.6. Correlation of motor task regressors with end-tidal CO_2_ regressor

To understand and quantify the potential confounding effect of task-correlated CO_2_ changes in our dataset, we visualized the interactions of the end-tidal CO_2_ regressor with the task regressors and calculated the correlations of an end-tidal CO_2_ regressor with the right, left, and right + left task regressors; two representative datasets with high correlation values are shown in **Figure 9**. In these datasets, increases in end-tidal CO_2_ are seen in the periods between tasks (**Figure 9A**). This observation is likely due to more shallow breathing during the hand grasp periods. The datasets were then modelled with and without the end-tidal CO_2_ regressor to assess the impact of accounting for the effects of CO_2_ variations during the scan. At the subject level, the addition of an end-tidal CO_2_ regressor reduced some regions with negative t-statistics and can be seen as regions of positive t-statistics related to the end-tidal CO_2_ regressor (**Figure 9B**). At the group level, a paired analysis was performed comparing these activation maps created with and without the end-tidal CO_2_ regressor in the model, and no significant clusters in motor regions were found (**Supplemental Figure 10**). The observed subject-level effects are likely attributed to the predominantly negative correlation of the end-tidal CO_2_ regressor with the task (**Figure 9C**). Across all scans, the correlation values varied widely, ranging from |r| = 0.063 to |r| = 0.64. Most correlation values between the end-tidal CO_2_ regressor and the task regressors (R, L, and R+L) were negative due to the increase in CO_2_ that occurred after each hand grasp ended.

**Figure 9.**
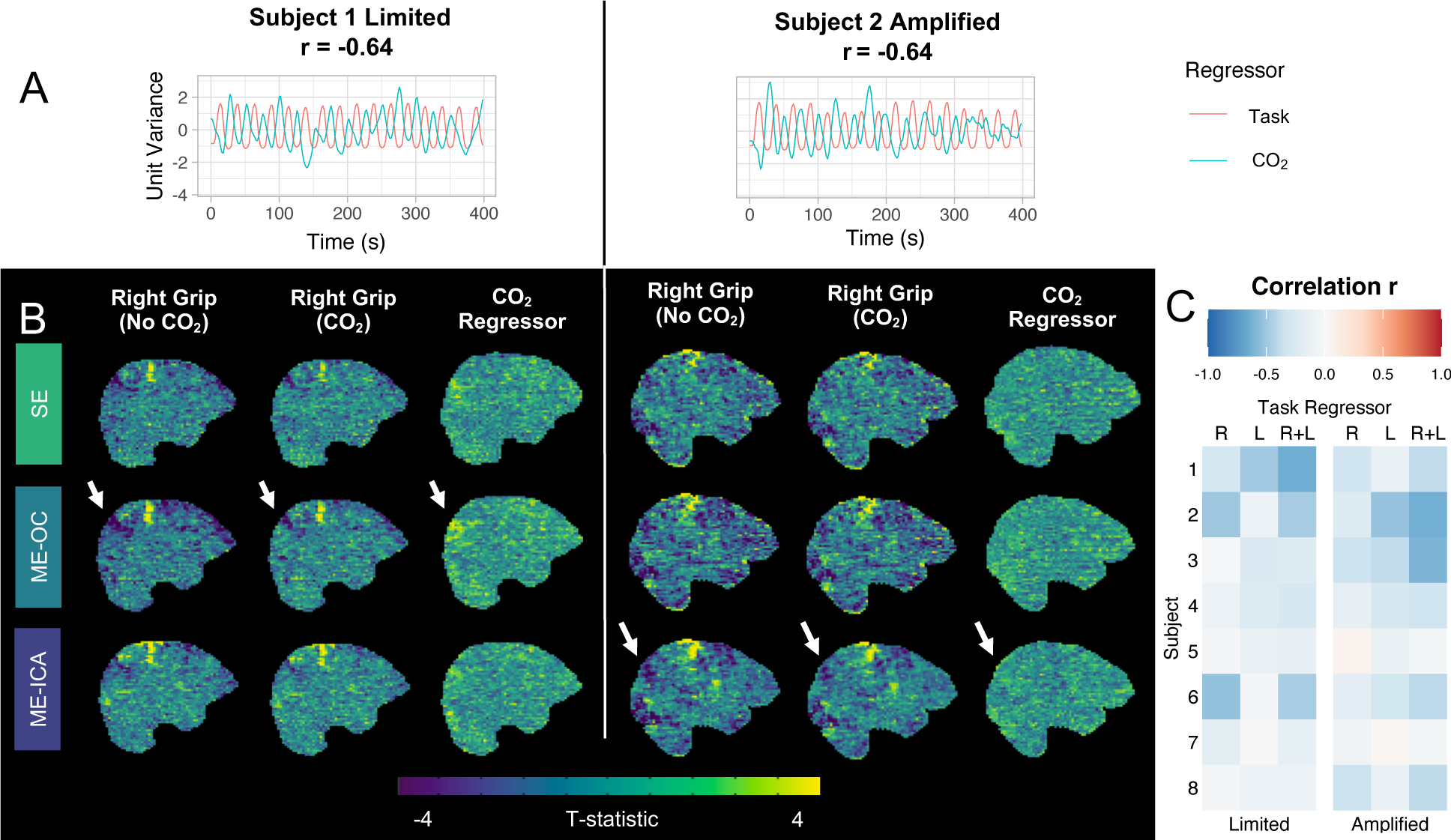
Two representative scans with high negative correlations of the end-tidal CO_2_ regressor with the summed task regressors are shown. (A) Timeseries of the end-tidal CO_2_ regressor and summed right and left task regressors, normalized to unit variance. End-tidal CO_2_ increases in the period between tasks. (B) Subject-level t-statistic maps from the right grip regressor are shown when running each model with and without an end-tidal CO_2_ regressor. T-statistic maps from the end-tidal CO_2_ regressor are also shown. Particularly in the ME-OC model, the addition of an end-tidal CO_2_ regressor reduced areas of negative t-statistics (indicated by arrows). This region shows a positive t-statistic in the end-tidal CO_2_ regressor map. (C) Correlations of the end-tidal CO_2_ regressor with the right (R), left (L), and summed (R+L) task regressors for each scan show primarily negative correlations across the dataset.

### 3.7. Group-level activation

Group-level activation maps were calculated for the Limited and Amplified motion conditions to determine the difference in the models’ abilities to identify primary regions of hand motor activity in datasets with low and high levels of head motion. Group activation maps for the three models are shown for the contrasts of Right Grip > 0 and Left Grip > 0 (**Figure 10**). In both Limited and Amplified motion conditions and with all three models, there are clusters in the hand-knob regions contralateral to the hand grasp. There are two expected areas of activation in the ipsilateral cerebellum, the V/VI and VIIIa/b lobules. The Right Grip > 0 Limited and Left Grip > 0 Amplified contrasts show significant clusters in both lobule regions, while the Right Grip > 0 Amplified and Left Grip > 0 Limited contrasts only show one cluster in one lobule region. The ME models result in slightly more robust activation in the cerebellum compared to SE, as demonstrated by larger clusters in the cerebellum, indicated by pink arrows in **Figure 10**.

**Figure 10.**
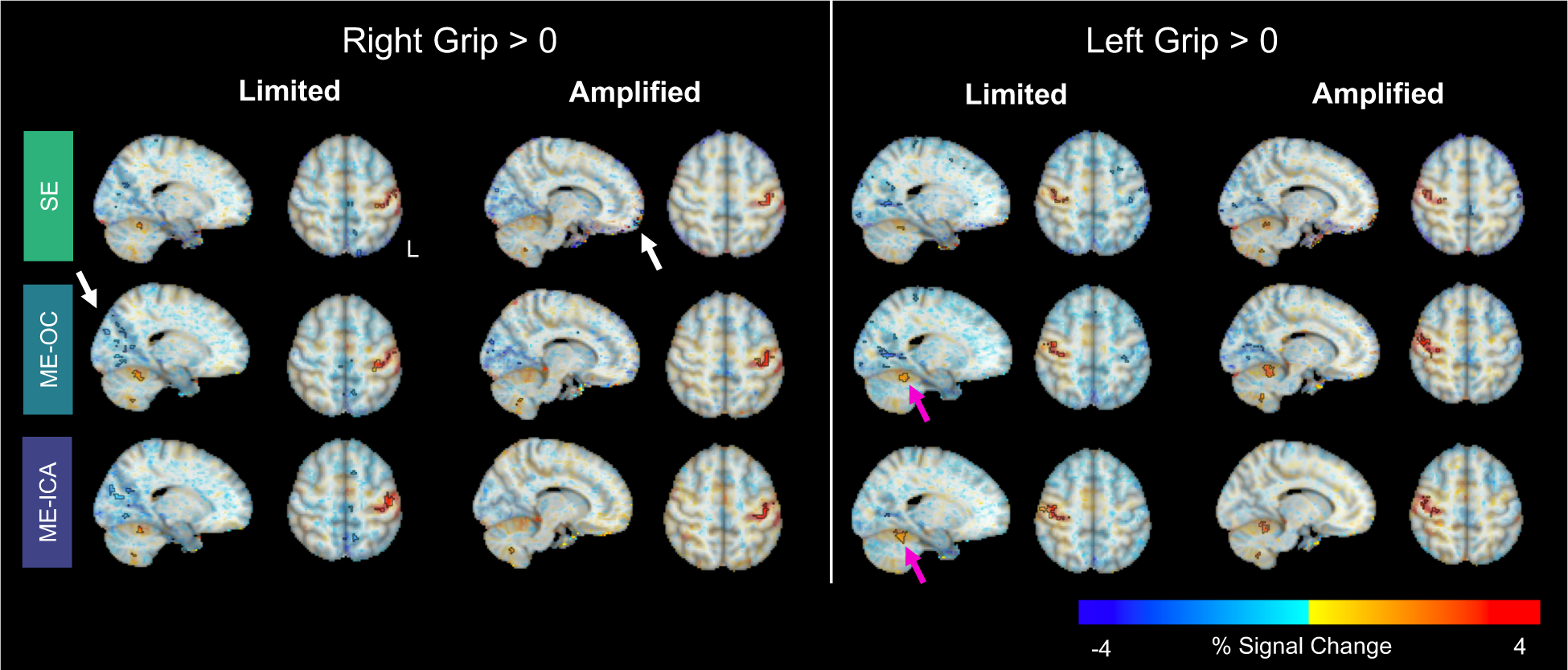
Group-level beta coefficient maps for the contrasts Right Grip > 0 and Left Grip > 0, shown for each analysis model and head motion condition. Opacity of beta coefficients is modulated by the t-statistic, as recommended by Taylor and colleagues (2023). Significant clusters are outlined in black, found by thresholding at p < 0.005 and clustering at α < 0.05. Pink arrows highlight cerebellar regions with robust activation in the ME models. White arrows highlight regions of negative clusters, which may be artifacts. Results that include the Subject 1 Amplified scan can be found in **Supplemental Figure 11**.

Other, primarily negative, clusters are seen in the occipital and frontal lobes and at the edges of the brain, which may be artifacts, highlighted with white arrows in **Figure 10**. The clusters in the inferior frontal lobes in the Amplified condition are reduced in the ME-OC and ME-ICA models compared to SE. The clusters in the occipital lobe are somewhat inconsistent across motion conditions and contrasts, with the ME-ICA model resulting in the lowest prevalence of occipital lobe clusters in all but the Right Grip > 0 contrast in the Limited condition. Quantitatively, paired group-level analyses between the Limited and Amplified motion conditions for each of the three models and grasp hands yielded no significant clusters in motor areas.

Quantitative comparisons of the group-level activation maps across the Limited and Amplified conditions and the three models for hand-knob and cerebellum motor ROIs are shown in **Figure 7B**. Similar to the subject-level results, group-level activation extent increases in the ME models compared to the SE model. The hand-knob and cerebellum regions in the ME models have greater median beta coefficients and t-statistics than the SE model.

## 4. Discussion

In this study, we tested ME-OC and ME-ICA against SE methods on a motor-task dataset with a range of task-correlated head motion amplitudes. In a population of healthy adults, we collected a set of scans with typical, limited head motion and a set of scans with amplified task-correlated head motion to simulate the challenges potentially experienced in a motor-impaired population. We examined the motion and breathing characteristics of these datasets to better understand the inherent artifacts and modeling challenges, and we showed our simulation was a reasonable approximation of motion challenges observed in stroke participants performing a similar task (**Figure 2**). Then, we applied SE, ME-OC, and ME-ICA models to identify activity in the brain associated with the hand grasp task. ME models demonstrated a better ability to dissociate head motion from the BOLD signal and reduce noise compared to SE. Additionally, ME-ICA models were better able to mitigate motion-related artifacts and provide potentially more stable and reliable activation estimates on a subject-level when task-correlated head motion was magnified. On the group level, all three models resulted in similar activation clusters in the motor areas of interest, with ME-OC and ME-ICA reducing some presumed artifactual negative clusters in other brain regions.

### 4.1. Magnitude and task-correlation of head motion are distinct phenomena

Through our simulation, we aimed to increase task-correlated head motion during a motor task; as expected, this simulation increased both average FD and the correlations of various directions of motion with the task (**Figure 1**). However, an increase in average FD was not always associated with a concurrent increase in task-correlation, and vice versa. For example, Subject 1’s Amplified scan had a much higher average FD than the other Amplified motion scans (FD = 1.85), but the degree of task correlation of the motion was similar. The same trend can be seen in the stroke datasets. Stroke Subject 1 had an average FD similar to that of the Healthy Amplified scans, as well as similar task-correlation in the Z direction. Stroke Subject 2 had much higher average FD, but lower task-correlations of motion. These observations suggest that two distinct, but related, complications may arise when dealing with increased head motion in motor-task fMRI: greater average head motion across the scan and greater task-correlation of head motion.

The analysis of potentially censored volumes showed that, using a threshold of FD > 0.5 that is considered ‘lenient’ by some studies (Power et al., 2014, 2017), 20-50% of volumes may be removed by censoring in our simulated “patient” data with Amplified movement (**Figure 2**). This level of data removal may be unacceptable in task-fMRI studies, and ME-ICA could be an important alternative to reduce strong motion-related signal changes and reduce loss of data.

### 4.2. ME-ICA mitigates motion-related artifacts in subject-level activation maps

To assess the ability of each model to reduce motion-related artifacts in the brain, we analyzed the relationship between DVARS and FD in each subject’s data and created gray plots for visualization of signal patterns across voxels. The relationship between DVARS and FD has been used in several previous studies to understand how well a model is able to dissociate the effects of head motion from changes in the BOLD signal (Kundu et al., 2013; Moia et al., 2021; Muschelli et al., 2014; Power, Silver, et al., 2019). We found that the ME-ICA model led to a significantly lower correlation between DVARS and FD compared to the SE and ME-OC models, particularly in the Amplified motion condition (**Figure 3**). Moia and colleagues (2021) observed the same trends across models when analyzing fMRI data during a breath-hold task, where task-correlated motion is also typically enhanced (Bright et al., 2009). Gray plots are another method of visualizing the timeseries of voxels across the brain to easily identify widespread patterns in fMRI data (Power, 2017; Power et al., 2018). Through gray plot visualization, ME-ICA was seen to reduce noise in white matter compared to SE and ME-OC, particularly in datasets with high amounts of head motion (**Figure 4**). This observation indicates a better ability of ME-ICA to reduce noise in regions not associated with the task.

We then observed how these differences in dissociation of head motion from BOLD signal manifested in subject-level activation maps. Subject-level activation maps are particularly important in populations that may exhibit subject-level vascular and neural impairment differences. For example, reliable subject-level activation maps can aid in analysis in a stroke population that stratifies participants by motor impairment level. Vascular differences in this population due to the stroke lesion may also make group-level analysis less feasible. We analyzed subject-level activation maps in the Limited and Amplified datasets to understand how the increased amplitude and task-correlation of head motion, as seen in motor-impaired populations, may influence our ability to reliably visualize motor activation.

In a scan with low average motion, we observed the primary effect of ME-OC and ME-ICA to be an apparent smoothing across the brain compared to SE (**Figure 5**). As average head motion increased in a scan, more artifacts became apparent in the SE and ME-OC conditions, primarily as noise at the edges of the brain and a banding artifact across the brain, possibly due to the interaction between the multi-band acquisition and head motion. ME-ICA largely mitigated these artifacts; Olafsson and colleagues (2015) also observed improvements when implementing ME-ICA with a simultaneous multi-slice protocol. The moderate, high, and very high motion examples shown in **Figure 5** are more representative of the level of motion seen in a motor-impaired population (**Figure 2**) and display the artifacts that might be encountered during a motor-task scan in these populations. At higher levels of motion, SE and ME-OC models may not sufficiently remove artifacts at the subject level, while ME-ICA models allow for cleaner subject-level maps.

Of note, a previous SE study also analyzed the effect of ICA on removing the effects of task-correlated motion during a motor task (Kochiyama et al., 2005). The authors found that their ICA approach led to fewer false negative errors than a traditional voxel-wise regression-based approach. However, they note a limitation of their method that reduces its efficacy when the head motion follows a similar block design to the task (i.e., movement at the beginning of the task, remaining at the same position, and a return to the original position at the end of the task). In our study, this was the type of movement we simulated in the Amplified condition, and we observed similar characteristics of movement in the Limited condition and in the Stroke Subject scans. Even with this type of problematic motion in our scans, we found the subject-level improvements described above; these improvements were potentially aided by the added benefits of ME acquisition and ME rho and kappa parameter information used to classify ME-ICA components. Different numbers of ICA components per dataset and specific implementation of these components in denoising methods may also vary results between studies that use ICA.

### 4.3. ME-ICA results in more stable and reliable beta coefficient estimates in the presence of large amounts of task-correlated head motion

We investigated the within-subject similarity of activation estimates by examining the voxel-wise spatial correlation between the Limited and Amplified motion scans for each subject. Spatial correlation has been used by several fMRI studies as a metric of similarity and stability across scans (Braban et al., 2023; Dipasquale et al., 2023; Gong et al., 2023; Kannurpatti et al., 2012; Yuan et al., 2013). The Limited motion scan for each subject served as an estimate of the ground-truth activation in our dataset. We found that ME-ICA demonstrated higher spatial correlations than SE when comparing both unthresholded maps and maps thresholded by significant positive activation (p_FDR_ < 0.05) to reduce effects of background noise. The Dice similarity coefficient was also higher when using ME-ICA compared to SE, when comparing areas of significant positive activation in the Limited and Amplified motion maps. These observations suggest that ME-ICA increases within-subject stability of beta coefficient estimates in scans with high task-correlated head motion. Similarly, Cohen and colleagues (2021) demonstrated that between-session spatial correlation increased when using ME-ICA, compared to SE data processed with ICA.

To investigate reliability of beta coefficient estimates, we also quantified motor activation in the subject-level maps through ROI analysis in regions associated with hand grasping: the hand-knob area of the motor cortex and cerebellum lobules V/VI and VIIIa/b. The association of hand grasp with activity in the hand knob of healthy participants has been well established (Cramer, Weisskoff, et al., 2002; Ehrsson et al., 2000; Keisker et al., 2009, 2010; Kuhtz-Buschbeck et al., 2008; Ludman et al., 1996; Thickbroom et al., 1998, 1999). The cerebellum lobules V/VI and VIIIa/b are associated with upper extremity activity and are also activated during hand and finger motor tasks (Ashida et al., 2019; Spencer et al., 2007; Stoodley, 2012). Therefore, we chose these regions to analyze the activation extent, median positive beta coefficient, and median positive t-statistic for each scan, comparing across models (**Figure 7A**).

The hand-knob ROIs were manually drawn for each participant to isolate the hand motor region. Since our Amplified motion condition involved voluntary head motion, we sought to exclude potential confounds of activity in the head motor area. The hand-knob region used in this study has been widely implemented by other groups to identify hand motor activity (Carlstedt et al., 2009; Cramer et al., 2001; Davare et al., 2008; Nakajima et al., 2020; Newton et al., 2005; Qiu et al., 2014). This definition of the hand motor area is also compatible with more recent understandings of the organization of the motor homunculus described by Gordon and colleagues (2023). This new homunculus model describes functional motor areas with distal body regions (e.g., hand) in the center and proximal body regions (e.g., shoulder) surrounding; this description still describes distal hand motor activity in one specific region of the motor cortex, in a comparable location to classical models of the homunculus. In the cerebellum, head motion is associated with activity in vermal lobule VI (Mottolese et al., 2013), which is an inter-hemispheric region that does not overlap with our defined motor cerebellum ROI.

Our hand-knob ROIs were drawn on each subject’s anatomical image, which had high spatial contrast to allow for proper delineation of the region. ROIs were then transformed to each subject’s functional space using a linear transformation (FLIRT). Since SE and ME analysis used different echoes for registration (second and first echo, respectively), the transformations may be slightly different. However, the echoes used in both SE and ME analyses had sufficient contrast to properly perform the registration steps.

In our study, we found activation extent in the selected hand-knob and cerebellum ROIs to be greater using the ME models compared to SE, likely due to increased sensitivity to activation aided by increased contrast-to-noise. Gonzalez-Castillo and colleagues (2016) had similar findings in their analysis of several types of task-fMRI data, showing a significant increase in activation extent from SE to ME models. The beta coefficient results varied across ROIs. In the cerebellum, ME models demonstrated a significantly lower median beta coefficient than the SE model. This result is also in line with the results of Gonzalez-Castillo and colleagues (2016), who observed a decrease in average effect size across different types of tasks.

Notably, our study found different trends for the beta coefficient in the hand-knob regions. With an ideal model, the beta coefficient would be similar between the Limited and Amplified conditions because the hand grasp was similar during both scans (**Figure 1A**) and therefore the magnitude of true BOLD activation should also be similar. The Limited condition scan, performed by young, healthy individuals, with head motion minimized by tactile feedback of tape on their foreheads, is our best approximation of ground truth activation for this dataset. We expected that a reliable measure of activation from the Amplified condition scans would have a beta coefficient more similar to the Limited condition scans. In particular, the hand-knob regions are heavily studied to measure hand motor activity, and accounting for the potential confounding effects of head motion there is critical. For example, the increase in hand-knob beta coefficient values in the Amplified compared to Limited conditions observed when using the ME-OC model may be an artifactual increase caused by the added head motion. The ME-ICA model has more comparable beta coefficients between motion conditions, suggesting it does a better job at representing the true activation in the motor cortex during the hand grasp when motion is high. A related finding was observed in a visual retinotopy task fMRI experiment performed by Steel and colleagues (2022), who also reported an increase in reliability of population receptive field parameter estimates when using ME-ICA compared to ME-OC, evaluated through correlation of parameter values across several datasets from the same subject.

We also examined the across-subject stability and consistency of beta coefficient estimates by analyzing the relationship of the median beta coefficient in each ROI with head motion. We found that ME-ICA leads to a lower relationship of the beta coefficient with both average FD and task-correlation of movement across all Limited and Amplified scans (**Figure 8**). This analysis provides additional evidence that SE and ME-OC models may artificially inflate beta coefficient estimates when task-correlated head motion increases, while ME-ICA aids in providing more stable estimates across scans with varied amounts of head motion.

Across both motion conditions, the median positive t-statistic increased in the ME models compared to SE, with no significant difference between ME-OC and ME-ICA. While Gonzalez-Castillo and colleagues (2016) similarly observed an increase in average t-statistic using ME models, they also found a significant increase from ME-OC to ME-ICA. As discussed above, in our study, the beta coefficient was lower in the ME-ICA versus ME-OC model during the Amplified condition. T-statistics increase with a larger beta coefficient and/or smaller model residuals; therefore, it is important to realize that the ME-ICA and ME-OC models had comparable t-statistics despite a lower ME-ICA beta coefficient. This indicates that the ME-ICA model likely had lower residuals of the model, in addition to a more reliable beta coefficient estimate in the hand-knob regions, during the Amplified condition. Although we cannot determine the accuracy of our activation estimates without complementary imaging modalities, these collective results suggest that ME-ICA is beneficial in our single-subject analysis of high-motion data.

### 4.4. In addition to head motion, end-tidal CO_2_ is also correlated with the motor task

Previous studies have found that task-correlated breathing may be an additional complication in motor-task fMRI, causing artifactual activation through variations in CO_2_ (Birn et al., 2006; Farthing et al., 2007) and motion (Power, Lynch, et al., 2019). Shallow breathing during the task can cause an increase in CO_2_, which leads to widespread vasodilation in the brain with varying timings (Bright et al., 2009; Chang et al., 2008). Although not the main focus of this study, we assessed the magnitude of this issue in our dataset by calculating the correlations between the end-tidal CO_2_ regressors and the hand grasp regressors. Correlations varied considerably across subjects and scans, with some correlations greater than |r| = 0.5 (**Figure 9C**). The direction of these correlations was primarily negative; end-tidal CO_2_ increased after the grasp periods (**Figure 9A**), likely due to shallower breaths during the task periods that lead to a gradual increase in arterial CO_2_ (Abbott et al., 2005).

Our approach to mitigating the confounds of task-correlated breathing was to add an end-tidal CO_2_ regressor to each model to account for the CO_2_ variation during the scan. To understand the effect of adding this regressor to our model, we analyzed our datasets with models that did not include the end-tidal CO_2_ regressor. On a subject level, the primary difference was that adding an end-tidal CO_2_ regressor reduced some areas of apparent negative activation (**Figure 9B**). As the end-tidal CO_2_ regressor was negatively correlated with the task regressors, this difference suggests that some variance in the data that was originally anti-correlated with the task regressors can be attributed to the end-tidal CO_2_ regressor after its addition to the model. On the group level, adding the end-tidal CO_2_ regressor to the model did not affect activation estimates in motor regions, suggesting that the spatial pattern of its effect is not consistent across participant and contribute to the fMRI signal more globally (**Supplemental Figure 9**). Future work focused on understanding these physiological confounds during motor tasks may consider using a lagged general linear model approach that has been used previously to account for spatially varying temporal alignment between the CO_2_ regressor and the voxel-wise BOLD fMRI timeseries (Moia et al., 2020, 2021; Stickland et al., 2021; Zvolanek et al., 2023). However, this exploration was outside the scope of this study, which focused on the use of ME-ICA to mitigate the effects of head motion.

Our findings emphasize the importance of recording and accounting for the effects of breathing during motor-task scans. Notably, non-motor tasks, such as working memory tasks, can similarly entrain breathing (Bright et al., 2020). If collecting CO_2_ data is not feasible, a respiration belt can also be used to measure breathing; a respiratory variance per time (RVT) regressor derived from respiration belt data has been shown to explain similar variance in the BOLD signal compared to an end-tidal CO_2_ regressor, when large breathing modulations are present (Zvolanek et al., 2023). In resting-state data, where breathing modulations are presumably smaller, RVT and end-tidal CO_2_ are potentially complementary in explaining BOLD signal changes (Chang & Glover, 2009; Golestani et al., 2015). Hence, collection of both CO_2_ and respiratory belt data may be useful depending on the study goals, magnitude of breathing variability, and feasibility of physiological signal acquisition.

### 4.5. Group-level activation maps are similar across models, even with increased head motion

Group-level analysis is commonly performed in motor-task fMRI studies to identify clusters of activation across subjects (Ehrsson et al., 2000; Keisker et al., 2009, 2010; Kuhtz-Buschbeck et al., 2008). To compare the three models’ performance in datasets with low and high motion, we did group-level analyses within the Limited and Amplified conditions (**Figure 10**). Unlike with the subject-level maps, there were no large visual differences between the conditions and models. All maps showed motor activation in the expected hand-knob and one or both of the cerebellum motor regions. In paired group-level analyses between the Limited and Amplified conditions for each of the three models, no significant differences in motor areas were found. The reason for the similarities between group-level results may be due to the variability in subject-level artifacts. **Figure 5** demonstrates the striping patterns that are the most obvious visual artifact in the high motion subject-level results. These striping artifacts vary in position and orientation between participants, while the primary area of motor cortex activation is largely similar. Hence, group-level analysis may average out these striping artifacts across participants, while retaining the common robust areas of cortical and cerebellar motor activation.

Lombardo and colleagues (2016) compared ME-ICA and ME-OC models during group-level task-fMRI analysis and found that effect size increased with ME-ICA; therefore, while we did not observe consistent differences at the group-level when using ME-ICA, this model may lead to group-level improvements in other instances. In this study, we found that SE models were robust to detecting motor activation in the motor cortex and cerebellum, even with datasets that have high motion. Consistent with the findings by Lombardo and colleagues, we found that ME models showed an increase in group-level beta coefficients and t-statistics across ROIs (**Figure 8B**). In most instances, ME-ICA also led to an additional increase compared to ME-OC.

While cortical and cerebellum motor regions were analyzed as ROIs in this study, other brain regions that are involved in sensorimotor processing may be important to consider in future studies. In particular, the thalamus has been implicated in motor control during fMRI studies of healthy and motor-impaired individuals (Charyasz et al., 2023; Errante et al., 2023; Mallol et al., 2007; Matsuda et al., 2009). In our study, we did not observe significant group-level clusters in the thalamus for either motion condition (Limited or Amplified), using any model (SE, ME-OC, ME-ICA), and therefore we did not include it in our ROI analyses. Studies that have demonstrated motor-related thalamus activity have done so during more dynamic tasks than the one performed in our study, for example, finger tapping and dynamic hand opening and closing (Charyasz et al., 2023; Matsuda et al., 2009). Since the thalamus has been implicated in the initiation and frequency of movement (Dacre et al., 2021; Lehéricy et al., 2006; MacMillan et al., 2004), the lack of significant thalamus clusters in our study may be due to our choice of the motor task. Our hand grasp paradigm did not involve several movement initiations, but instead one movement initiation followed by a maintenance of force produced.

Given our observations, if a study aims to detect results at the group level, SE fMRI data may be sufficient to map motor activation during a simple motor task even in high motion datasets. However, in clinical populations that may exhibit high task-correlated motion, subject-level analysis is particularly important: clinical research questions and patient diagnosis may motivate achieving insight into single-subject neural activity patterns, and it may not even be feasible or appropriate to implement group analyses because of pathological differences in anatomy and vascular physiology between subjects. In these cases, our study demonstrates that ME-ICA is critical to disentangling the effects of head motion from true motor activation, resulting in more reliable subject-level conclusions.

### 4.6. Limitations

Our motion simulation used an instructed head nod to amplify the task-correlated head motion during the scans. However, this is not a perfect substitute for the increased head motion exhibited by clinical populations. The added voluntary head motion in our Amplified scans may add a *neural* confound to the motor activation analysis. To limit our sensitivity to the neural effects of the voluntary head motion, we manually drew a hand-knob ROI to isolate the region of the motor cortex involved in hand motion from the region involved in head motion. We also confirmed that the motion characteristics of our Amplified scans were similar to those of two scans from individuals with chronic stroke (**Figure 2**).

Another challenge is identifying a ground truth in our study. While there are expected areas of activation in the hand knob and cerebellum motor areas that we used as our a priori ROIs for analysis (J. Diedrichsen et al., 2011; Jörn Diedrichsen et al., 2009; Yousry et al., 1997), we did not have a clear ground truth for the magnitude of the beta coefficient and activation extent within these ROIs. The Limited motion condition in our study involved young, healthy individuals performing a task with low levels of motion; this was our best approximation of accurate beta coefficient measurements using fMRI methods. Electroencephalography (EEG) may be another method to determine beta coefficient accuracy in the cortical hand knob areas, though it would not be able to investigate subcortical cerebellum motor areas.

Additionally, ME-ICA is unlikely to salvage datasets with very large amounts of head motion. In our study, the Amplified motion dataset approximated the motion characteristics of Stroke Subject 1. However, Stroke Subject 2 exhibited much higher head motion, and we have not thoroughly tested whether ME-ICA would be effective in mitigating the effects of this extreme degree of head motion. However, our findings in our outlier with very high motion (Subject 1 Amplified dataset) suggest that ME-ICA may demonstrate similar improvements in datasets with FD up to 1.8 (**Figure 5**). Future work should probe the limits of ME-ICA in dealing with large amounts of head motion.

### 4.7. Considerations for future studies

The hand grasping task studied here can be associated with high levels of task-correlated head motion, but other motor tasks may have lower associated head motion. For example, commonly used finger tapping tasks involve more isolated and limited movements and likely will not encounter the same challenges with head motion. However, tasks such as hand grasping, and more proximal or complicated upper extremity movements, are important in the study of different neurological pathways and clinical conditions such as stroke (Ellis et al., 2016; McPherson et al., 2018; Wilkins et al., 2020). Also, individuals with motor impairments might not be able to perform finger individuation tasks, but could perform hand grasping or other less precise movements for the study of brain motor activity.

Other external methods of minimizing head motion may be possible, such as using a customized head mold (Power, Silver, et al., 2019) or applying tape across the participant’s forehead to add tactile feedback of head motion (Krause et al., 2019). In this study, we implemented the addition of tape across the forehead to further reduce head motion during our Limited motion condition and allow for a better approximation of ground truth activation. Additionally, real-time motion tracking for prospective motion correction techniques can be implemented to reduce false activations (Maclaren et al., 2013; Schulz et al., 2014). Each of the previously described techniques can potentially be used in combination with ME-ICA to further minimize motion artifacts.

Although there are clear benefits to ME acquisition and ME-ICA analysis, there are also certain limitations. ME acquisition requires longer TRs in order to collect multiple echoes, which may not be feasible or desirable in every application. Manual classification of ICA components for ME-ICA analysis also takes some training for consistent classification across scans, and additional hands-on analysis time to perform the classification. This manual step may introduce potential for unwanted bias; therefore, we pre-determined a list of classification rules (*2.2.3*) to limit the variation in component classification and have publicly shared the classifications used in this study to aid reproducibility (Reddy et al., 2023). For large datasets, in which it may not be feasible to manually classify components, it may be possible to explore creation of a user-defined decision tree using new and developing features of tedana (Ahmed et al., 2023; Dupre et al., 2021).

## 5. Conclusion

Motor-task fMRI is prone to task-correlated head motion that confounds activation results, particularly in certain clinical populations. In this study, we collected an fMRI dataset from a healthy population who performed a hand grasp task with Limited and Amplified task-correlated head motion to simulate a motor-impaired population. We analyzed these data using three models: single-echo (SE), multi-echo optimally combined (ME-OC), and multi-echo independent component analysis (ME-ICA). On the subject level, ME models better dissociated the effects of head motion from the BOLD signal and led to increased t-statistics in brain motor regions. In scans with high levels of motion, ME-ICA additionally mitigated artifacts and led to more reliable beta coefficient estimates. Overall, we show that ME-ICA is a useful tool for analyzing motor-task data with high levels of task-correlated head motion, which is particularly important in studies of clinical populations where subject-level analysis is the goal.

## CRediT author contribution statement

**Neha A. Reddy:** Conceptualization, Methodology, Software, Formal analysis, Investigation, Data curation, Writing – original draft, Writing – review & editing, Visualization, Project administration. **Kristina M. Zvolanek:** Investigation, Writing – review & editing. **Stefano Moia:** Methodology, Writing – review & editing. **César Caballero-Gaudes:** Conceptualization, Methodology, Writing – review & editing. **Molly G. Bright:** Conceptualization, Methodology, Writing – review & editing, Supervision, Project administration, Funding acquisition.

## Acknowledgements

N.A.R. and K.M.Z. were supported by the National Institutes of Health under a training program (T32EB025766). K.M.Z was supported by the National Heart, Lung, and Blood Institute of the National Institutes of Health under Award Number F31HL166079. The content is solely the responsibility of the authors and does not necessarily represent the official views of the National Institutes of Health. This work was supported by the Center for Translational Imaging at Northwestern University. This research was also supported by the Spanish Ministry of Economy and Competitiveness (Ramon y Cajal Fellowship, RYC-2017-21845), the Basque Government (BERC 2018-2021 and PIBA_2019_104), and the Spanish Ministry of Science, Innovation and Universities (MICINN; PID2019-105520GB-100).

