## Supplemental Material for "Denoising task-correlated head motion from motor-task fMRI data with multi-echo ICA"

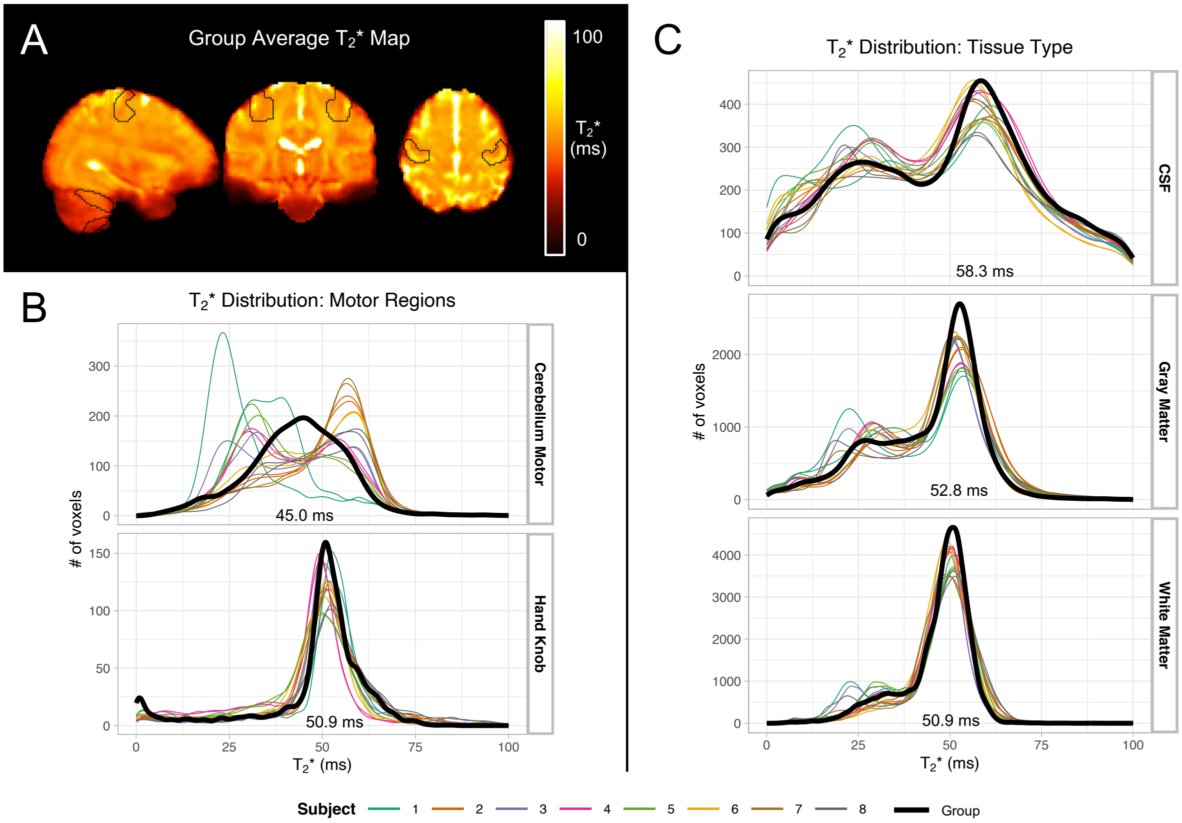


**Supplemental Figure 1**. (A) $T_2$* map averaged across all subjects and scans. Subject-level $T_2$* maps were transformed to standard space, then averaged. Hand knob and cerebellum motor region masks used in ROI analysis are outlined in black. (B) Distribution of $T_2$* in hand knob and cerebellum motor region masks, shown for individual subject scans and group average map. $T_2$* values at peak density in each region are listed. (C) Distribution of $T_2$* in CSF, gray matter, and white matter regions, shown for individual subject scans and group average map. Tissue region masks were created by segmenting the MNI brain with FSL FAST, then thresholded at 0.75. $T_2$* values of the group average map at peak density in each region are listed.

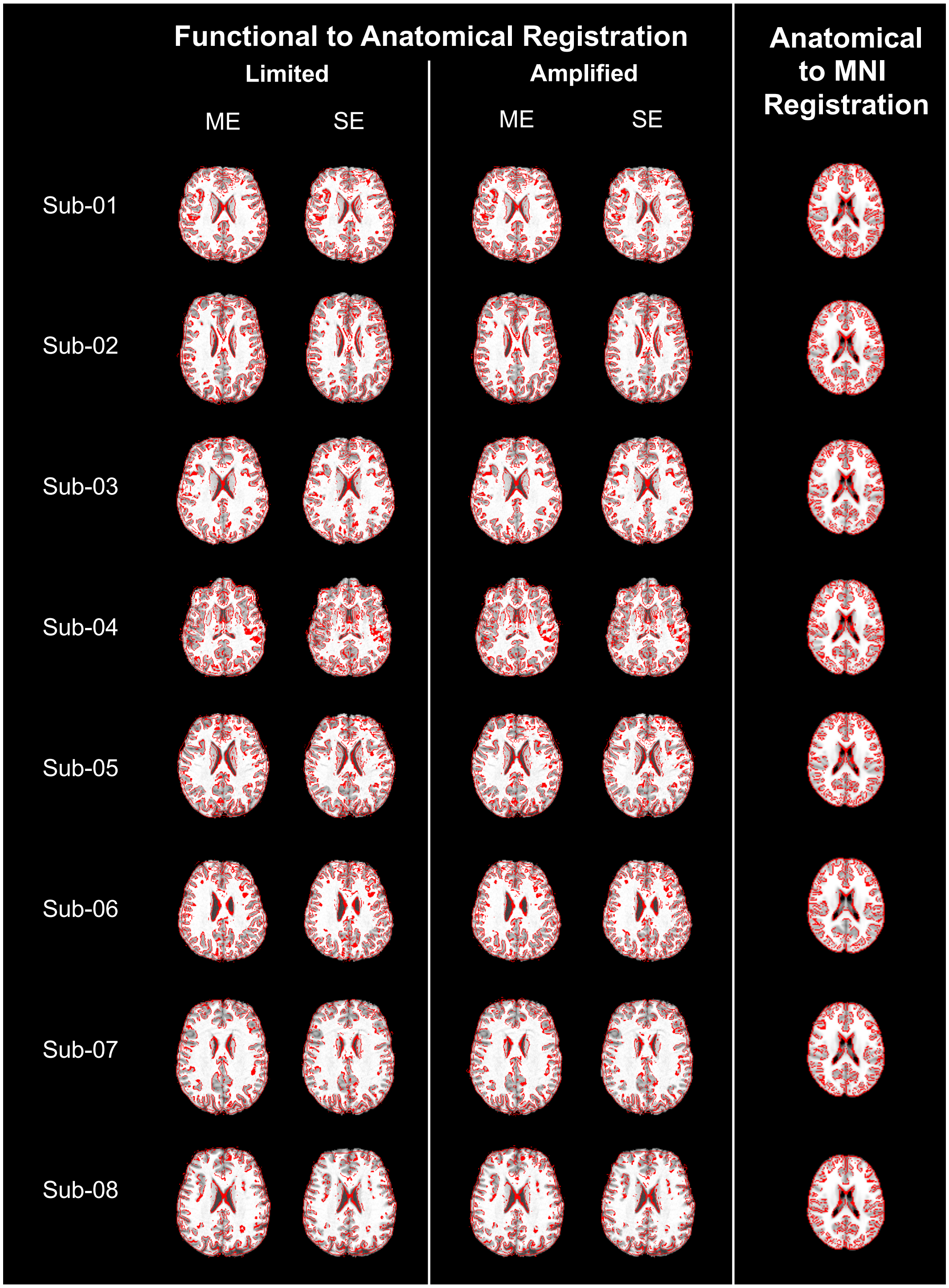


**Supplemental Figure 2**. Demonstration of all registration steps performed in this study, with slices chosen to include lateral ventricle. Each registration step was visually inspected. For functional to anatomical registration, the edges of the registered functional single band reference image were overlaid on the anatomical image using FSL's slicer. The alignment of the lateral ventricles and gray matter-white matter borders was examined. Functional to anatomical registration was checked for each functional scan (Limited and Amplified), and separately for multi-echo (using first echo) and single-echo (using second echo). Similarly, for anatomical to

MNI registration, the edges of the registered anatomical image were overlaid on the MNI image and examined. All registrations were determined to be satisfactory based on the visible alignment of anatomical landmarks.

**Supplemental Table 1.** Scan parameters for stroke participant functional scans. Parameters for Stroke Subject 2 fully match those of the healthy participant datasets.

| **Subject** | **Scan** | **TR (s)** | **TE (ms)** | **FA (°)** | **MB factor** | **GRAPPA** | **Voxel size ($mm^3$)** |
|---|---|---|---|---|---|---|---|
| Stroke 1 | 1 | 1.3 | 34.4 | 62 | 4 | None | 2x2x2 |
| Stroke 1 | 2 | 2 | 20 | 90 | None | 2 | 2x2x2 |
| Stroke 2 | 1 | 2 | 10.8/28.03/45.26/62.49/79.72 | 70 | 4 | 2 | 2.5x2.5x2 |
| Stroke 2 | 2 | 2 | 10.8/28.03/45.26/62.49/79.72 | 70 | 4 | 2 | 2.5x2.5x2 |

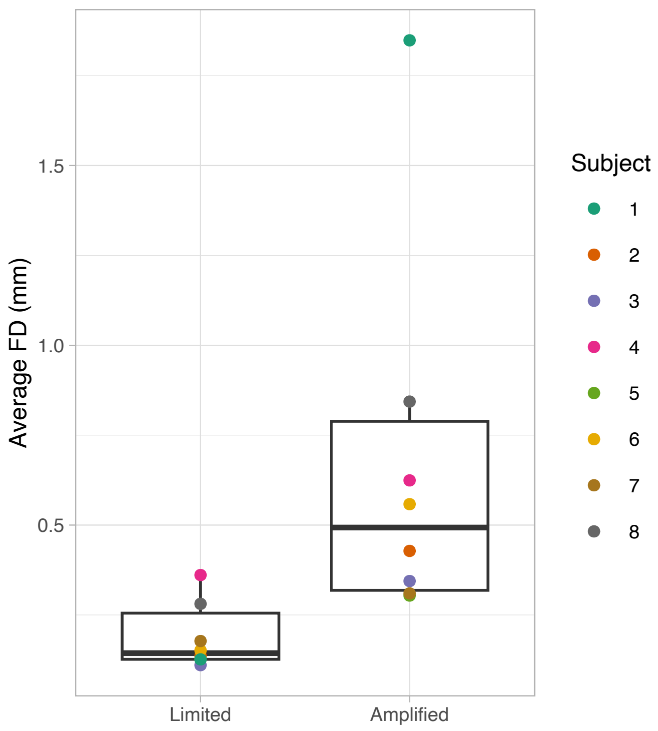


**Supplemental Figure 3.** Average Framewise Displacement (FD) for each subject and scan. The Amplified condition scan from Subject 1 was excluded from spatial correlation, ROI, and group activation analysis as an outlier because its average head motion (FD) was greater than the third quartile for FD + 1.5 times the interquartile range.

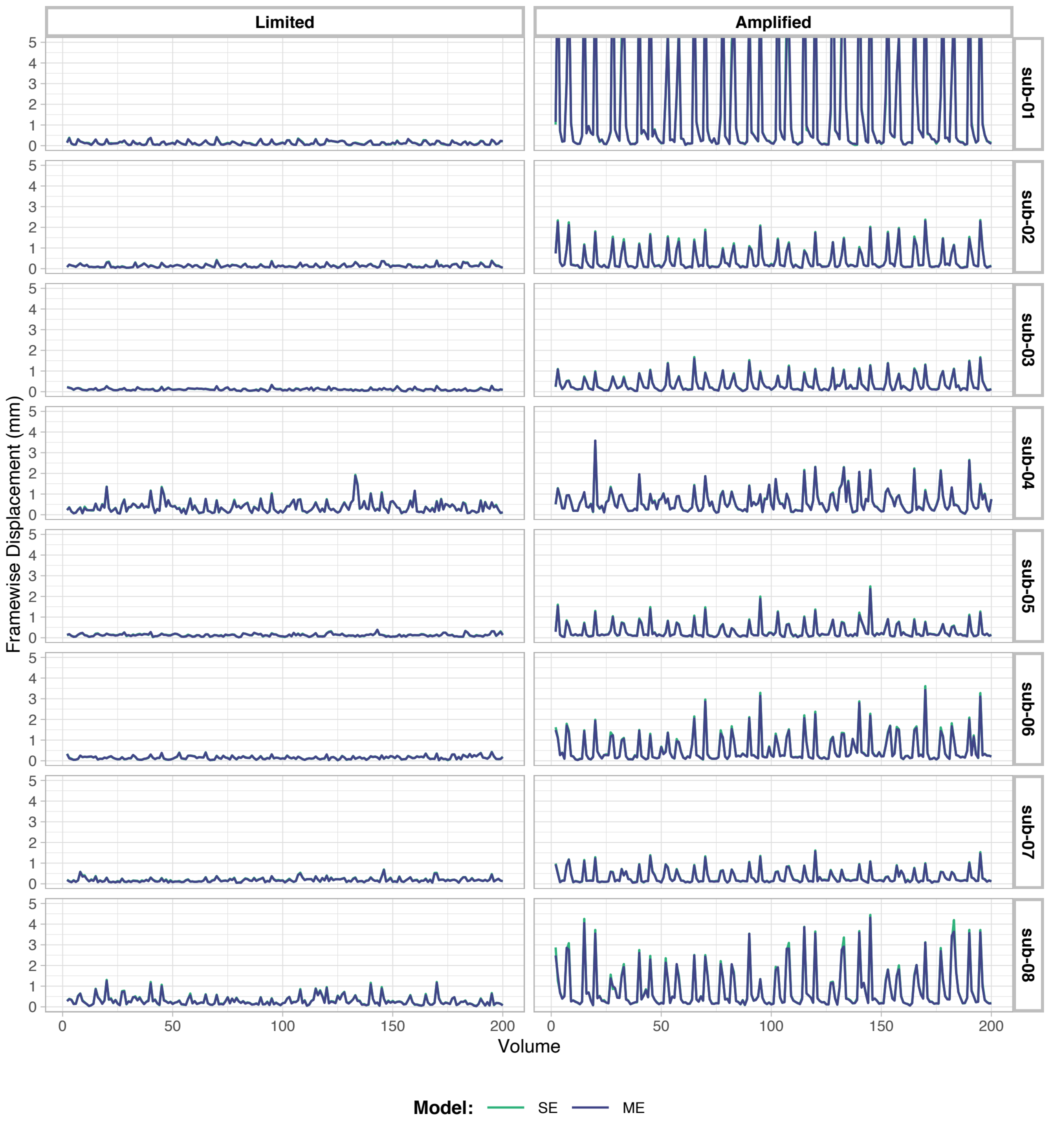


**Supplemental Figure 4.** Framewise Displacement (FD) values across each scan for SE analysis (volume realignment parameters obtained from the second echo) and ME-OC and ME-ICA analysis (volume realignment parameters obtained from the first echo). SE FD peaks are slightly higher for sub-06 and sub-08 in the Amplified condition. Note that the sub-01 Amplified scan was excluded from group analysis due to high levels of motion. Notice that the traces of FD are nearly identical regardless of using the first or second echo to estimate the realignment parameters.

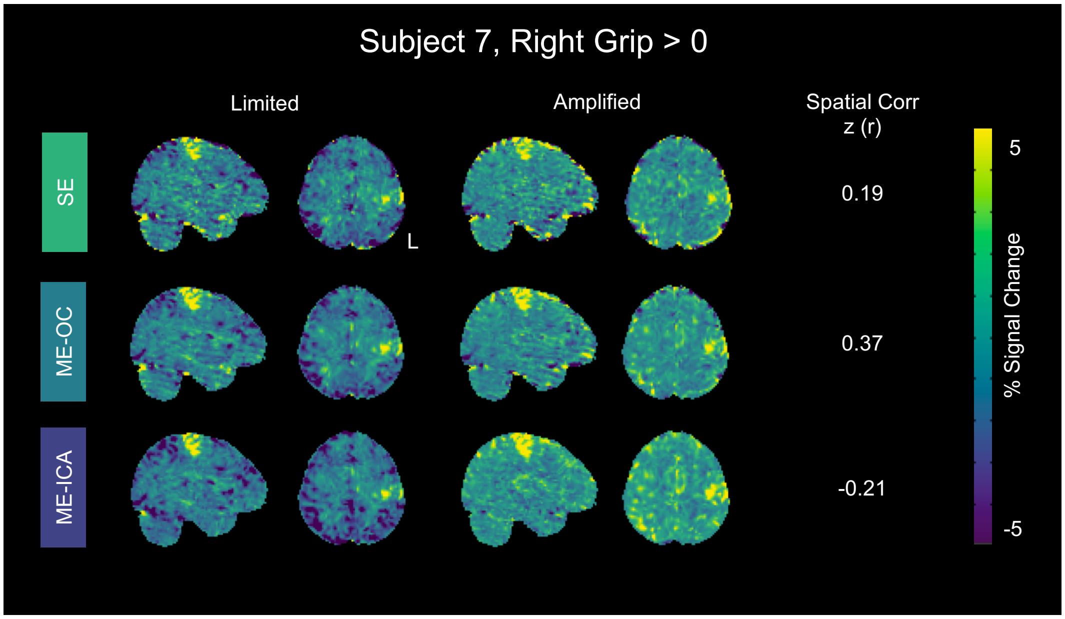


**Supplemental Figure 5.** Subject 7 maps in MNI space of percent BOLD signal change related to the right grip task performed with Limited and Amplified motion. Pearson r correlation values shown were calculated between unthresholded maps, then converted to Fisher z. All models show regions of negative activation in the Limited motion map that are positive in the Amplified motion map. This effect is most prominent with the ME-ICA model, leading to a negative spatial correlation. However, the regions in question are outside motor areas and have relatively low t-statistics that do not pass thresholding at $p_{FDR} < 0.05$.

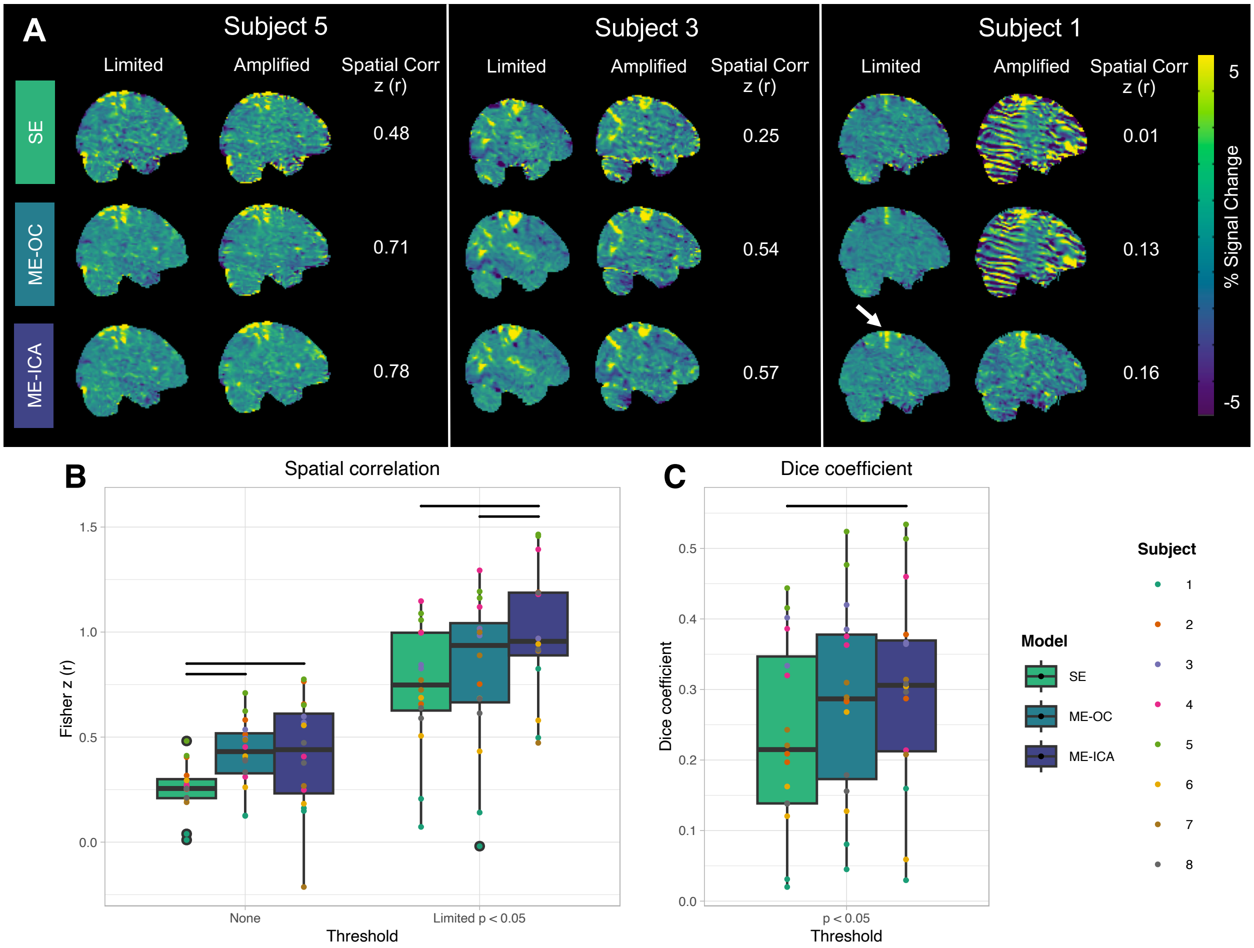


**Supplemental Figure 6. Same figure as Figure 6 in the main text but panels (B) and (C) include Subject 1**. (A) Subject-level maps in MNI space of percent BOLD signal change related to the right grip task performed with Limited and Amplified motion. Three representative scans with high, moderate, and low levels of spatial correlation between Limited and Amplified maps are shown. Pearson r correlation values shown were calculated between unthresholded maps, then converted to Fisher z. The arrow indicates the expected region of activation in the motor cortex. (B) Spatial correlation between Limited and Amplified maps. Correlation was calculated between unthresholded maps and between maps that were thresholded by voxels with significant positive activation in the Limited map. Significant positive activation was determined by thresholding the Limited map at $p_{FDR} < 0.05$, using a t-statistic accounting for degrees of freedom in the model. Only areas of positive activation were retained. A mask of these significant positive voxels was applied to Limited and Amplified beta parameter maps before correlation was calculated. Maps were thresholded to show correlation after reducing effects of background noise voxels on spatial correlation estimates. Subject-level maps in MNI space from RightGrip and LeftGrip activation were included. (C) Dice similarity coefficients were calculated between areas of significant positive activation in Limited and Amplified scans, respectively. Significant voxels were found as in (B), independently calculated for Limited and Amplified scans. Subject-level maps in MNI space from RightGrip and LeftGrip activation were included. Dots represent values from individual scans. Bars indicate significant difference between models ($p < 0.05$, Bonferroni-corrected).

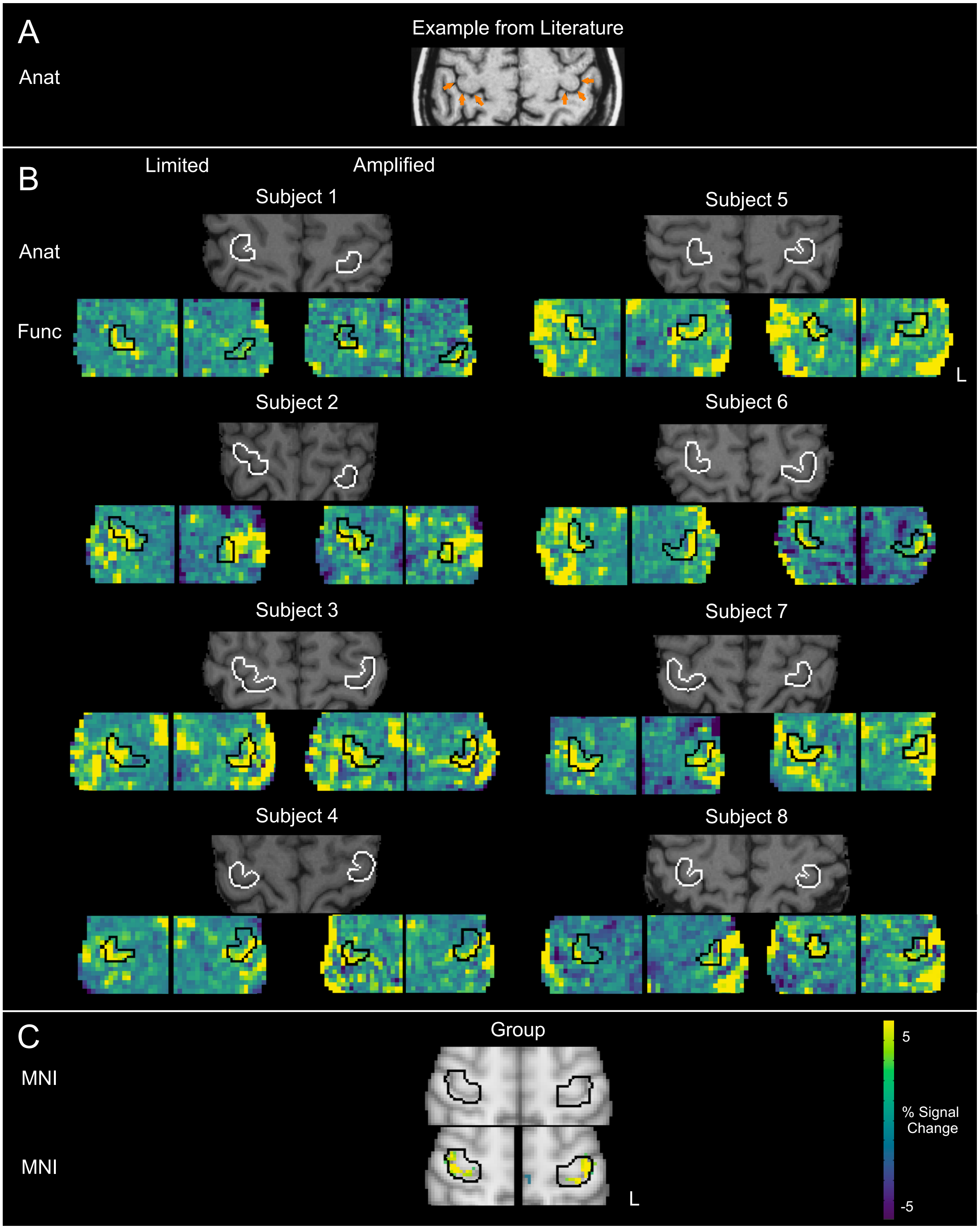


**Supplemental Figure 7.** Hand knob ROIs. (A) Example hand-knob ROI in anatomical space (Anat) from Yousry and colleagues (1997), used as a guide for manual identification of hand-knob ROIs in this study. 10% of hemispheres were reported to follow an 'epsilon' shape instead of the more common ‘inverted omega’ shown here. (B) Hand-knob ROIs in subject space. For each subject, masks of the right and left hand-knob regions were manually drawn. Masks were drawn in anatomical space (Anat), then the region that aligned with gray matter was retained. This mask was then dilated by two voxels before transformation to subject functional space for use in analysis (Func). For each subject, the final anatomical mask is shown (Anat). The left hand-knob mask is shown with the right grasp beta coefficient maps of the Limited and Amplified condition scans analyzed with the ME-ICA model, for demonstrative purposes. The right hand-knob mask is shown on top of the left grasp beta coefficient maps. Axial slices shown here were chosen to highlight the primary areas of overlap with motor activation in the Limited condition scan, then approximately the same slices were chosen for the Amplified condition scan. (C) Hand-knob ROIs in MNI space used in group analysis. Masks of the right and left hand-knob regions were drawn in MNI space, then dilated by two voxels. Shown here are the hand-knob masks in MNI space. The left hand-knob mask is also shown with the right grasp significant clusters at the group-level of the Limited condition scans analyzed with the ME-ICA model. The right hand-knob mask is shown on top of the left grasp significant clusters.

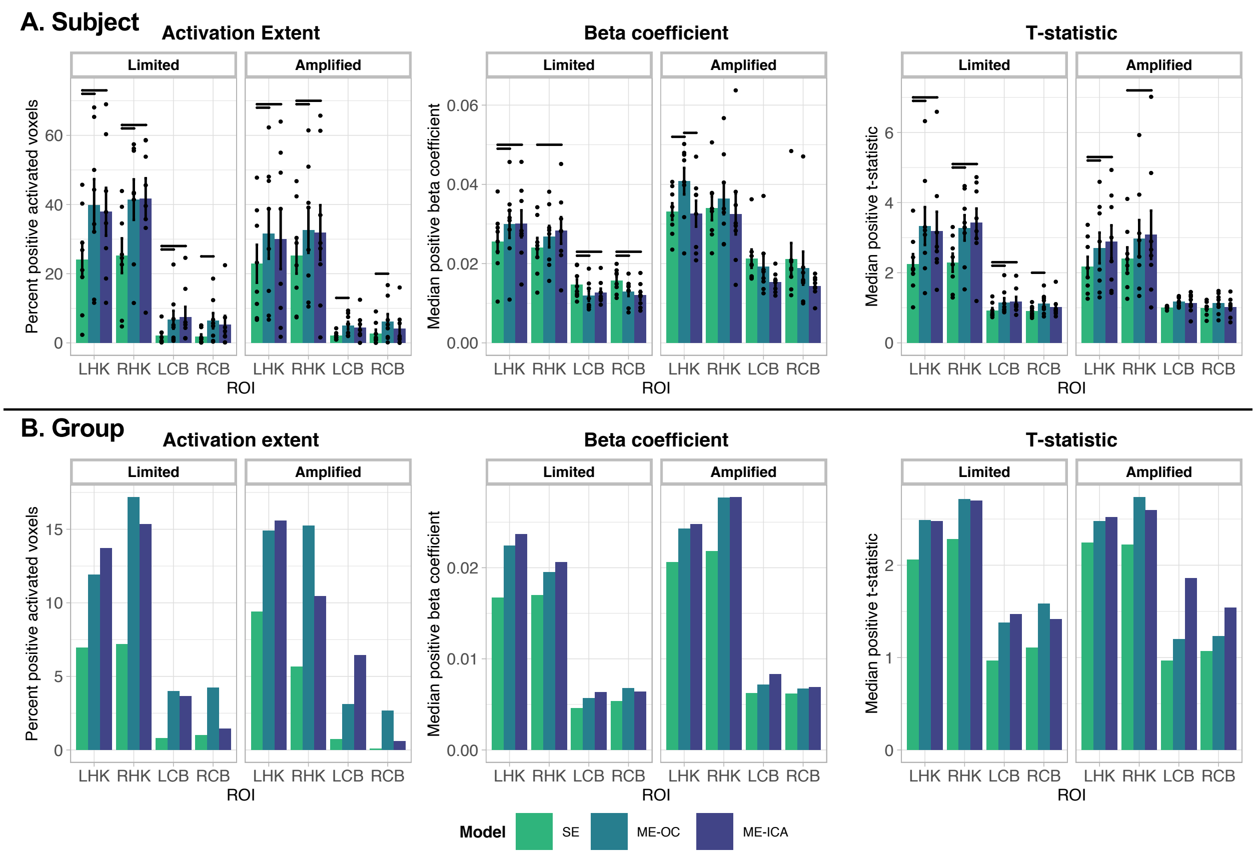


**Supplemental Figure 8. Same figure as Figure 7 in the main text but Amplified group includes Subject 1**. (A) Subject-level activation extent, beta coefficient, and t-statistic values for each scan associated with response to hand grasp tasks. Dots represent individual subject values. Vertical bars indicate standard error across subjects. Horizontal bars indicate significant difference between models ($p < 0.05$, Bonferroni-corrected). (B) Group-level activation extent for clustered group activation maps, and beta coefficient and t-statistic values for unthresholded group activation maps associated with response to hand grasp tasks. In (A) and (B), values are shown for four ROIs: values associated with right-hand grip in the left-hand knob (LHK) and right cerebellum hand motor areas (RCB) and values associated with left-hand grip in the right-hand knob (RHK) and left cerebellum hand motor areas (LCB).

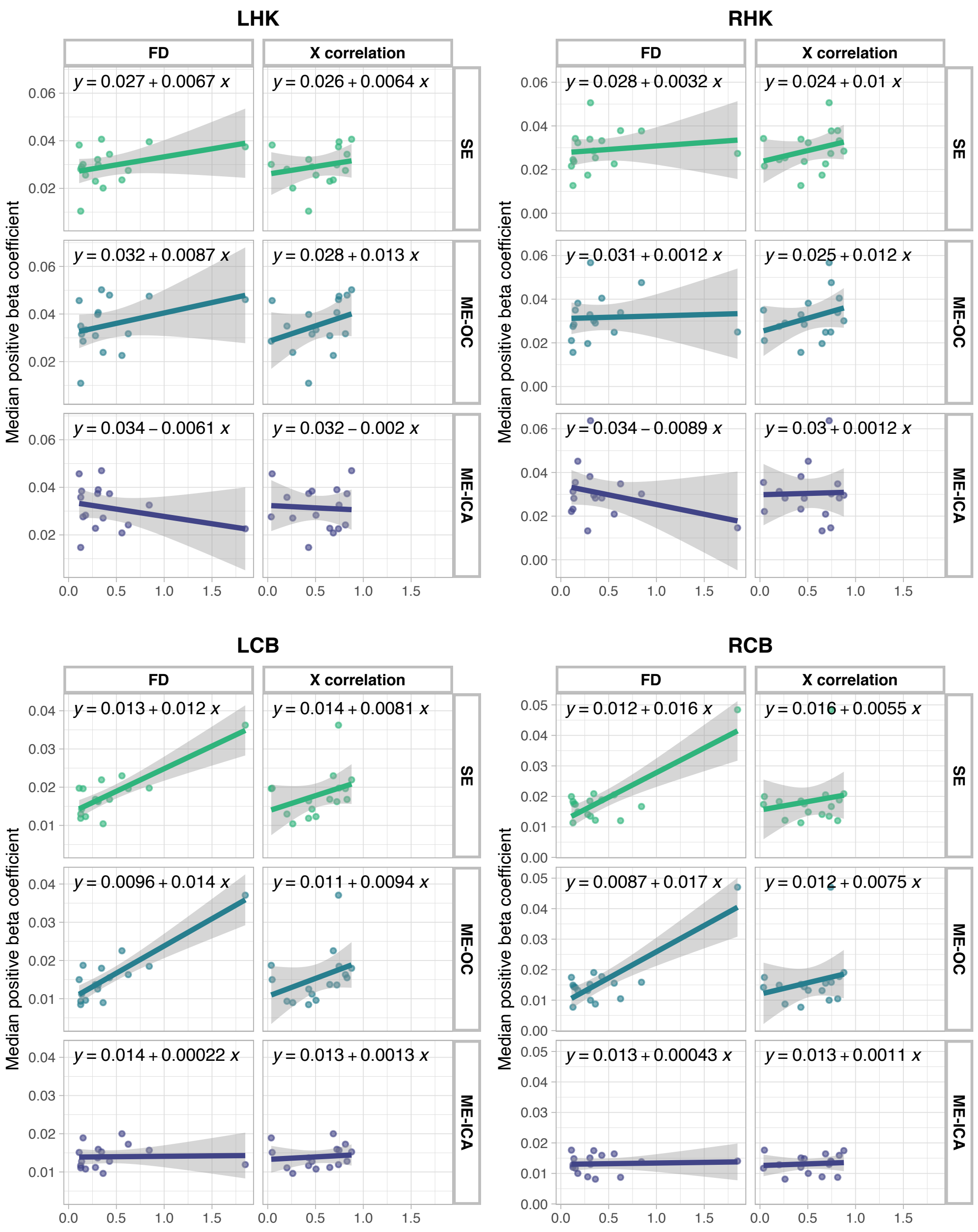


**Supplemental Figure 9. Same figure as Figure 8 in the main text but results include Subject 1**. Median beta coefficients calculated using SE, ME-OC, and ME-ICA models shown for each Limited and Amplified motion scan, plotted against the average FD and against the task correlation with motion in the X direction for each scan. The Amplified motion dataset from Subject 1 is included in these results. Beta coefficient values are shown for four ROIs: values associated with right-hand grip in the left hand knob (LHK) and right cerebellum hand motor areas (RCB) and values associated with left-hand grip in the right hand knob (RHK) and left cerebellum hand motor areas (LCB). For all but one case (RHK, FD correlation), ME-ICA results in the lowest relationship (slope) of the beta coefficient with average motion and task-correlation of motion.

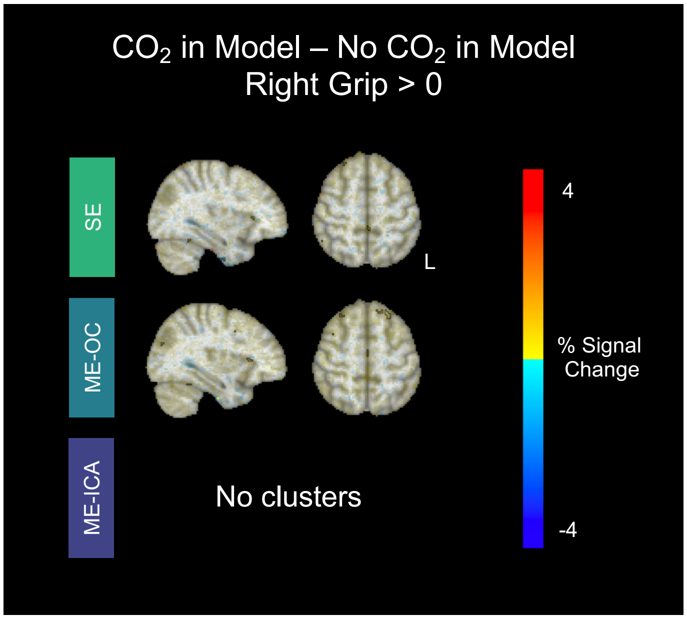


**Supplemental Figure 10.** Results of a paired group-level analysis of Right Grip activation during the Limited motion task, comparing the activation maps created *with* the end-tidal $CO_2$ regressor in the model to the activation maps created *without* the end-tidal $CO_2$ regressor in the model. Paired group analysis was performed with 3dttest++ (AFNI), then results were thresholded at $p < 0.005$ and clustered at $\alpha < 0.05$, using a bi-sided t-test (3dFWHMx, 3dClustSim, 3dClusterize, AFNI). For the SE and ME-OC models, a few small clusters were found across the cortex in regions not associated with the motor task. For the ME-ICA model, no clusters were found. This suggests that the spatial pattern of the $CO_2$ effect on the motor activation maps may not be consistent across participants.

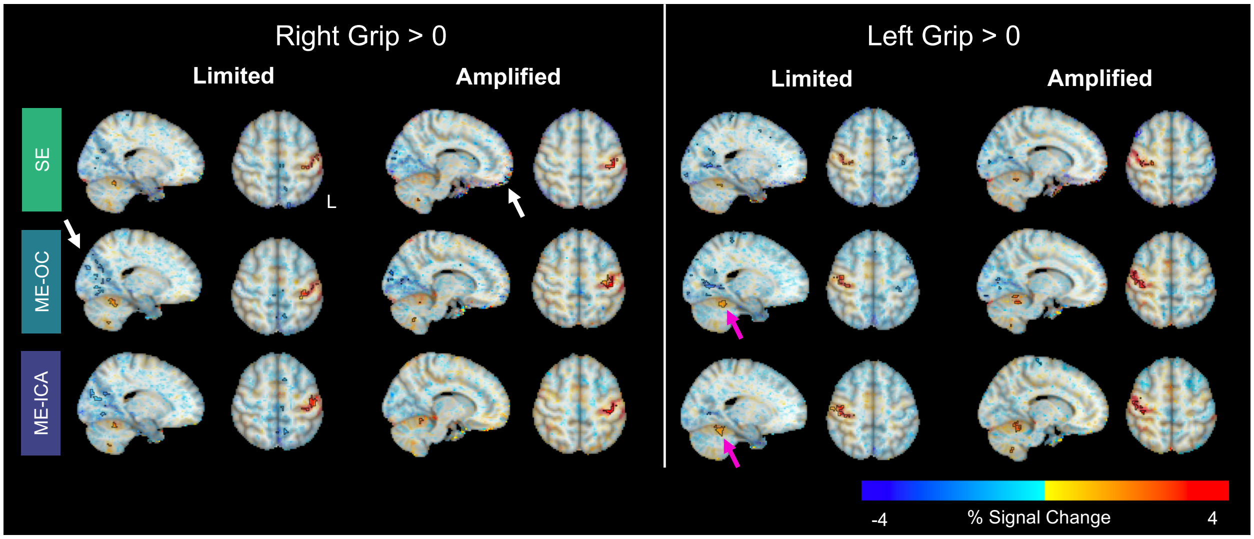


**Supplemental Figure 11. Same figure as Figure 10 in the main text but Amplified group includes Subject 1**. Group-level beta coefficient maps for the contrasts Right Grip > 0 and Left Grip > 0, shown for each analysis model and head motion condition. Opacity of beta coefficients is modulated by the t-statistic, as recommended by Taylor and colleagues (2023). Significant clusters are outlined in black, found by thresholding at $p < 0.005$ and clustering at $\alpha < 0.05$. Pink arrows highlight cerebellar regions with robust activation in the ME models. White arrows highlight regions of negative clusters, which may be artifacts.
